# Modeling phylogenetic biome shifts on a planet with a past

**DOI:** 10.1101/832527

**Authors:** Michael J. Landis, Erika J. Edwards, Michael J. Donoghue

## Abstract

The spatial distribution of biomes has changed considerably over deep time, so the geographical opportunity for an evolutionary lineage to shift into a new biome may depend on how the availability and connectivity of biomes has varied temporally. To better understand how lineages shift between biomes in space and time, we developed a phylogenetic biome shift model in which each lineage shifts between biomes and disperses between regions at rates that depend on the lineage’s biome affinity and location relative to the spatiotemporal distribution of biomes at any given time. To study the behavior of the biome shift model in an empirical setting, we developed a literature-based representation of paleobiome structure for three mesic forest biomes, six regions, and eight time strata, ranging from the Late Cretaceous (100 Ma) through the present. We then fitted the model to a time-calibrated phylogeny of 119 *Viburnum* species to compare how the results responded to various realistic or unrealistic assumptions about paleobiome structure.

Ancestral biome estimates that account for paleobiome dynamics reconstructed a warm temperate (or tropical) origin of *Viburnum*, which is consistent with previous fossil-based estimates of ancestral biomes. Imposing unrealistic paleobiome distributions led to ancestral biome estimates that eliminated support for tropical origins, and instead inflated support for cold temperate ancestry throughout the warmer Paleocene and Eocene. The biome shift model we describe is applicable to the study of evolutionary systems beyond *Viburnum*, and the core mechanisms of our model are extensible to the design of richer phylogenetic models of historical biogeography and/or lineage diversification. We conclude that biome shift models that account for dynamic geographical opportunities are important for inferring ancestral biomes that are compatible with our understanding of Earth history.

## INTRODUCTION

Biomes are ecologically and climatically distinct species assemblages that vary in size, shape, and continuity across geographical regions, in large part due to regional differences in temperature, precipitation, seasonality, altitude, soil types, and continentality (Whittaker 1970; Wolfe 1985; Olson et al. 2001; Mucina 2019). The diversity of biomes occupied by particular lineages also varies considerably, with some clades exhibiting strict associations with particular biomes, and others showing multiple transitions between biomes over time (Donoghue and Edwards 2014). Although it is accepted that clade-wide variation in regional biome occupancy was generated and is maintained by evolutionary forces including speciation, extinction, dispersal, and adaptation to new biomes, it remains difficult to estimate exactly when, where, and under what conditions phylogenetic lineages first shifted into the biomes that their descendants inhabit today.

In current practice, ancestral regions and biome affinities are often estimated independently of one another, and then relationships between regions and biomes are compared post hoc (e.g., Crisp et al. 2009; Weeks et al. 2014). Although such studies yield important evolutionary insights, the estimates themselves do not account for how lineages might move between regions or adapt to newly encountered biomes given the temporally variable spatial configuration of biomes across regions. Conceptually, how a biome is geographically distributed should influence how easily a lineage might disperse into a new region or shift into a new biome, an effect Donoghue and Edwards (2014) termed geographical opportunity. One strategy to model the effect of geographical opportunity first defines discrete regions that are exactly coincident with modern day biomes, and then assumes that species within a given region occur within the corresponding biome. Cardillo et al. (2017) carried out such an analysis in studying the biogeography of the Australian plant clade, *Hakea* (Protaceae), using method features developed by Matzke (2014), where total regional area and shared perimeter lengths tuned dispersal rates between bioregions. This innovative strategy depends crucially on the uniformity of biome composition within each region. Larger, discrete regions may very well be dominated by a single biome type, yet still be composed of assorted dominant, subdominant, and marginal biome types at local scales.

More importantly for our purposes, defining geographical opportunity based on modern biome features (such as area and shared perimeter), may be problematic in instances where the spatial distribution of biomes has changed considerably over time, since those changes should also influence when and where ancestral lineages shift between regions and/or biomes. For example, if woodlands dominated a particular region until the rise of grasslands, that might inform when a grassland-adapted lineage first dispersed into that region. That is, if the presences or absences of biomes among regions influence modern species ranges, then temporal variation in regional biome availability should similarly influence our models of range evolution.

To model how paleoecological dynamics might influence range evolution, Meseguer et al. (2015) fitted ecological niche models (ENMs) to fossil data so as to limit the connectivity between regions for biogeographical models that estimate ancestral ranges (Ree and Smith 2008). While this strategy is quite promising, its current form requires that the clade under study (*Hypericum* of Hypericaceae, in their case) has a sufficiently rich fossil record over space and time to inform the ENM. It also assumes that all lineages face the same, broad ecological limitations to range evolution, independent of what particular biome affinity each lineage possesses at a given moment. Although the quality of the fossil record is largely out of our control, the second assumption could be relaxed: ideally, if a clade contains sub-lineages that specialize in both woodland and in grassland habitats, any particular lineage’s range should be principally limited by the availability of the specific biome to which that lineage is adapted, rather than being constrained based on a broader, clade-wide average of grassland and woodland lineages.

In this paper, we aim to address the aforementioned challenges facing current phylogenetic models of biome shifting by incorporating four key properties: (1) that biome shifts and dispersal events share a common state space over biomes and regions, (2) that any discrete region may contain a number of different biomes, (3) that the geographical structure of biomes within and between regions can vary over time, and (4) that lineages adapted to different biomes and located in different regions will experience different dispersal rates between regions and different shift rates into new biomes. We begin by introducing a graph-based approach to characterize the availability, prevalence, and connectivity of regional biomes through time, building on the framework introduced by Landis (2017). We then develop an event-based evolutionary process using a time-stratified continuous-time Markov chain that models biome shifts and dispersal given the ways in which biome distributions have changed over time. Because the exact influence of extrinsic geographical factors and/or ecological structure is bound to vary from clade to clade, the degree of influence of such features on the evolutionary model are treated as free parameters to be estimated from the data itself.

To explore the possible importance of paleobiome structure on lineage movements among biomes, we apply our model to *Viburnum*, a clade of ∼165 species that originated in the Late Cretaceous and are today found in tropical, warm temperate, and cold temperate forests throughout Eurasia and the New World. We generated paleobiome graphs for these three mesic forest biomes across six continental regions for eight major time intervals over the past hundred million years. Fitting the model to our *Viburnum* dataset all-but-eliminates the possibility of a cold temperate origin of the clade. This is consistent with our understanding of the important biogeographic role of a “boreotropical” flora (see below) during the Paleocene and Eocene, and with our recent fossil-based ancestral biome estimates in *Viburnum* (Landis et al. 2020).

## METHODS

### Viburnum phylogeny and biogeography

*Viburnum* (Adoxaceae) is a clade of ∼165 extant plant species that originated just before the Cretaceous-Paleogene (K-Pg) boundary, roughly 70 Ma. Previous studies of phylogenetic relationships (Clement et al. 2014; Spriggs et al. 2015; Eaton et al. 2017) and divergence times (Spriggs et al. 2015; Landis et al. 2020) provide a firm basis for understanding the order and timing of lineage diversification events in *Viburnum*. In this study, we focus on a subsample of 119 *Viburnum* species with relationships that are highly supported by phylogenomic data (Eaton et al. 2017; Landis et al. 2020) and whose divergence times were time-calibrated under the fossilized-birth death process (Heath et al. 2014) as described in Landis et al. (2020).

*Viburnum* species occupy six continental-scale regions: Southeast Asia, including the Indoaustralian Archipelago and the Indian subcontinent; East Asia, including China, Taiwan, and Japan; Europe, including the North African coast, portions of the Middle East, and the Azores and the Canary Islands; a North American region north of Mexico; a Central American region that includes Mexico, Cuba, and Jamaica; and the South American Andes. The choice of these biogeographic regions is discussed in more detail in Landis et al. (2020). Briefly, these regions were selected because: (1) each is an area of *Viburnum* species/clade endemism; (2) there are no individual *Viburnum* species that occur in more than one of these regions; and (3) there are significant barriers to the migration of viburnums between these regions today and in the relevant past. As a concrete example, consider our recognition of North America and of Mexico, Central America, and the Caribbean as two distinct biogeographic regions for *Viburnum*. Multiple *Viburnum* species are endemic to the forests in each of these regions, and none of these species spans between them. Importantly, for mesic forest plants such as *Viburnum*, these two regions are entirely separated from one another by an extensive zone of low-lying arid lands in southeastern Texas and northeastern Mexico, with vegetation generally classified as xeric shrubland or mesquite-chaparral savanna. These areas are uninhabitable by viburnums today, and probably have not been accessible to these plants since widespread drying and the spread of C4 grasses commenced by at least the late Miocene (e.g., Godfrey et al. 2018). Using our criteria, the most difficult division is between the southeast Asian and eastern Asian regions. Although these do both harbor endemic species/clades, and these species are adapted to different environmental conditions, there are now, and have been in the past, opportunities for north-south migration between mesic forests of the two regions.

Finally, we note that our biogeographic regions are not strictly defined in terms of tectonic plates as they do not determine the movements and distributions of *Viburnum*. Thus, based on the criteria above, we split the vast Eurasian plate into a European region and two Asian regions. Likewise, the boundary between the North American and the Caribbean plates appears to have had no impact on *Viburnum* movements as there are individual species (e.g., *V. hartwegii*) that span the two.

Across those regions, living viburnums are affiliated with mesic forest biomes and show widespread parallel evolution of leaf form, leafing habit, and physiology coincident with transitions between warmer and colder biomes (Schmerler et al. 2012; Chatelet et al. 2013; Spriggs et al. 2015; Scoffoni et al. 2016; Edwards et al. 2017). Five extinct *Viburnum* lineages are known by their fossil pollen grains recovered from North American and European locales. Four of these fossils are older samples (48 to 33 Ma) from paleofloral communities that we previously judged to be warm temperate or subtropical (Landis et al. 2020). For our analyses in this study, we defined three mesic forest biomes based on annual temperatures and rainfall patterns (Edwards et al. 2017). Tropical forests have high temperatures and precipitation year-round, showing little seasonality. Warm temperate forests, which include paratropical, lucidophyllous, and cloud forests, vary seasonally in temperature and precipitation, but do not experience prolonged freezing temperatures during the coldest months. Cold temperate forests also experience seasonal temperatures and precipitation, but average minimum temperatures drop below freezing in at least one of the coldest months.

Because we are interested in how biome states and regional states evolve in tandem, we constructed a set of 3 × 6 = 18 compound states that we call biome-region states. Throughout the paper, we identify the biome-region state for a lineage in biome state *i* and region state *k* with the notation (*i,k*). However, in practice, we encode biome-region states as integers with values from 1 to 18. Biome-region state codings for *Viburnum* are translated from Landis et al. (2020), though here we combine cloud forests and warm temperate forests into a single warm temperate category.

All of the 119 Viburnum species we included in our analysis are found in a single biome, except for two East Asian (*V._chinshanense* and *V._congestum*) and one North American (*V. rufidulum*) species, which reside in forests possessing both warm and cold temperate elements. Those three species were coded as ambiguous for the relevant biome-region states. While it may be more accurate to describe those species as occupying multiple biomes, the model that we will soon define assumes that lineages occupy only one biome-region state at any given time. The biome-region character matrix and the time-calibrated phylogeny for *Viburnum* that we used are hosted on DataDryad (https://doi.org/10.5061/dryad.pvmcvdnhj).

### Model overview

Our aim is to model a regional biome shift process that allows changes in the spatiotemporal distribution of biomes to influence the likelihood of a lineage shifting between biomes and dispersing between regions. This process can be described in terms of interactions between two fundamental subprocesses: the biome shift process and the dispersal process.

The biome shift process models when and where lineages shift into new biome types. The probability of a biome shift clearly depends on intrinsic and extrinsic factors that govern how readily a lineage might adapt to the conditions in a new biome, which involves a myriad of factors that we do not fully explore here. Rather, we focus specifically on modeling the effect of geographical opportunity on biome shifts (Donoghue and Edwards 2014). For example, it is plausible that a species inhabiting the warm temperate forests of Europe might have shifted into the tropical biome during the Early Eocene, a period when tropical rain forests could be found at latitudes as extreme as 60^∘^ N. In contrast, a biome shift within Europe from a warm temperate to a tropical biome would be less likely today or during any time after the global cooling trend that began with the Oligocene.

The dispersal process models how lineages move between regions. The rate of dispersal between regions should depend on how connected those regions are for a given biome affinity. Returning to the Europe example, a tropical lineage in Southeast Asia might, all else being equal, have a relatively high dispersal rate into Europe during the Early Eocene, when Europe was predominantly tropical and warm temperate, as compared to today, when Europe is dominated by temperate and boreal forests. This rate would, however, also be influenced by geographical connectivities at the time. In this case, the existence of the Paratethys Sea and the Turgai Strait may have influenced plant migrations at different latitudes during the Eocene, although the fragmented northern shore of the Tethys Sea facilitated the movement of tropical and subtropical plants throughout much of that period (Tiffney 1985a, b; Tiffney and Manchester 2001).

How the biome shift and dispersal processes may behave in response to an evolving biome structure is depicted in Figure 1. By characterizing known features of paleobiome structure (Fig. 1A) into adjacency matrices (Fig. 1B), we can differentiate between probable and improbable phylogenetic histories of biome shifts and dispersal events (Fig. 1C) based on time-dependent and paleobiome-informed biome shift rates (Fig. 1D) and dispersal rates (Fig. 1E). Of the two regional biome shift histories in Figure 1C, the first history invokes three events that are fully congruent with the underlying paleobiome structure. The second history requires only two events, yet those events are incongruent with the paleobiome structure. But which regional biome shift history is more probable? Assigning probabilities to histories must depend not only on the phylogenetic placement and age of the regional biome shift events, but also on the degree to which the clade evolves in a paleobiome-dependent manner. We later return to how this unknown behavior of the evolutionary process may be estimated from phylogenetic data, only after we define a probabilistic model for the process.

**Figure 1.**
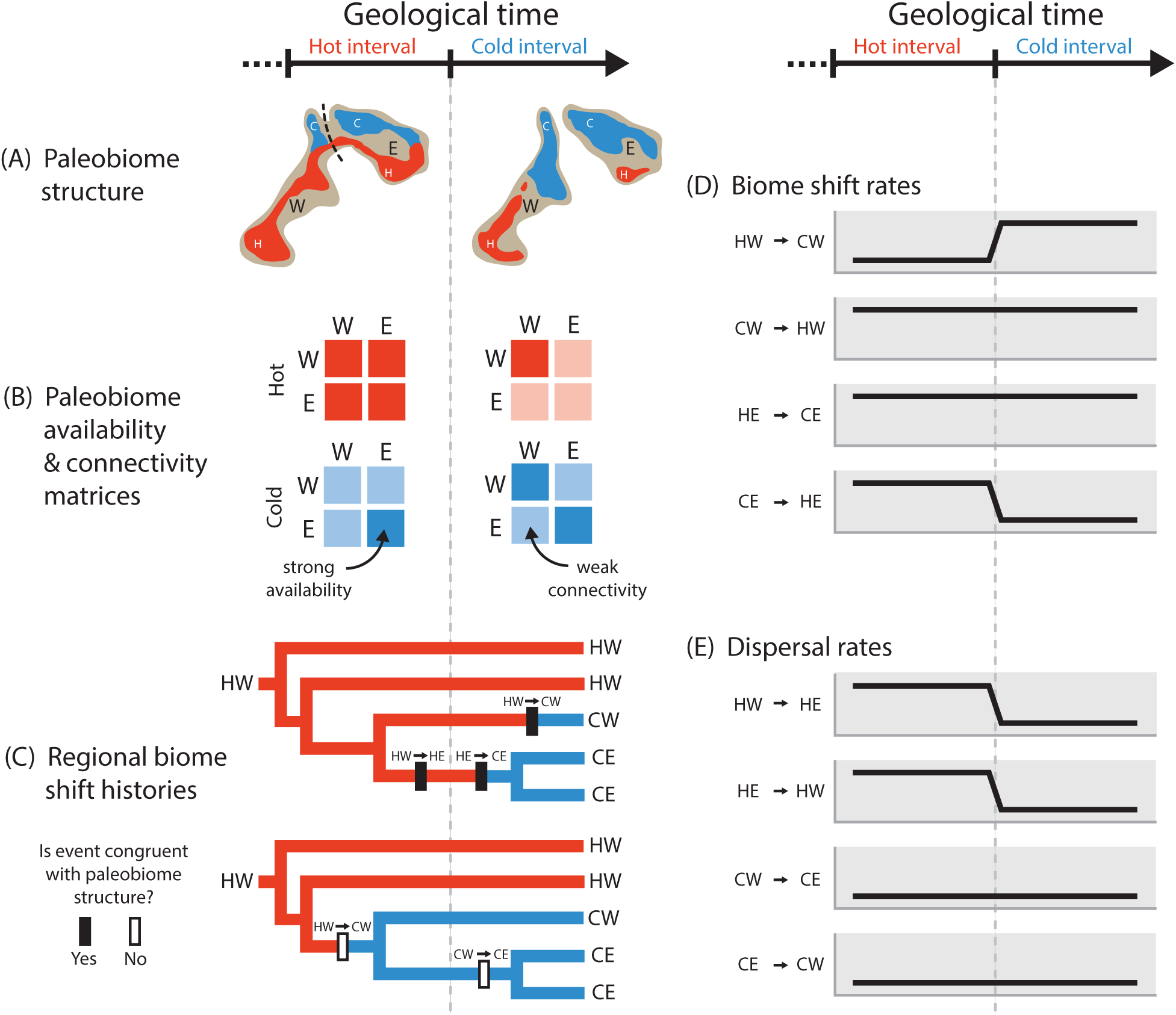
Cartoon of the relationship between paleobiome structure and a regional biome shift process. The left and right panels are aligned to the same geological time scale that is divided into a Hot (red) interval followed by a Cold (blue) interval. (A) Maps of paleobiome structure with two regions, East (E) and West (W), and two focal biomes of interest, Hot (H) and Cold (C), in which the expansive Hot biome is replaced by the Cold biome as the East and West regions separate. (B) Paleobiome adjacency matrices encode the availability of biomes within regions and the connectivity of biomes between regions based on whether paleobiome features are strong (dark) or weak (light). Diagonal elements reflect biome availability within regions while off-diagonal elements report biome connectivity between regions. (C) Two possible regional biome shift histories for a phylogeny with a western, hot-adapted (HW) origin. Lineages shift between biomes at rates that depend on the availability of biomes within the lineage’s current region and disperse between regions at rates that depend on connectivity of the lineage’s current biome between regions. The two histories require (top) or do not require (bottom) evolutionary events to be congruent with paleobiome structure. (D) Time-dependent biome shift rates for the four possible events: HW to CW, CW to HW, HE to CE, and CE to HE. (E) Time-dependent dispersal rates for the four possible events: HW to HE, HE to HW, CW to CE, and CE to CW.

### An evolving spatial distribution of biomes through time

Biome availability and connectivity has evolved over time. We summarize these dynamics with a series of time-dependent graphs that are informed by the paleobiological and paleogeographical literature (Fig. 2). To define our paleobiome graphs, we consulted global biome reconstructions generated by Wolfe (1985), Morley (2000), Graham (2011, 2018), Fine and Ree (2006), Jetz and Fine (2012), and Willis and McElwain (2014) which we then corroborated with biome reconstructions quantitatively estimated using the BIOME4 model (Prentice et al. 1992; Kaplan et al. 2003).

**Figure 2.**
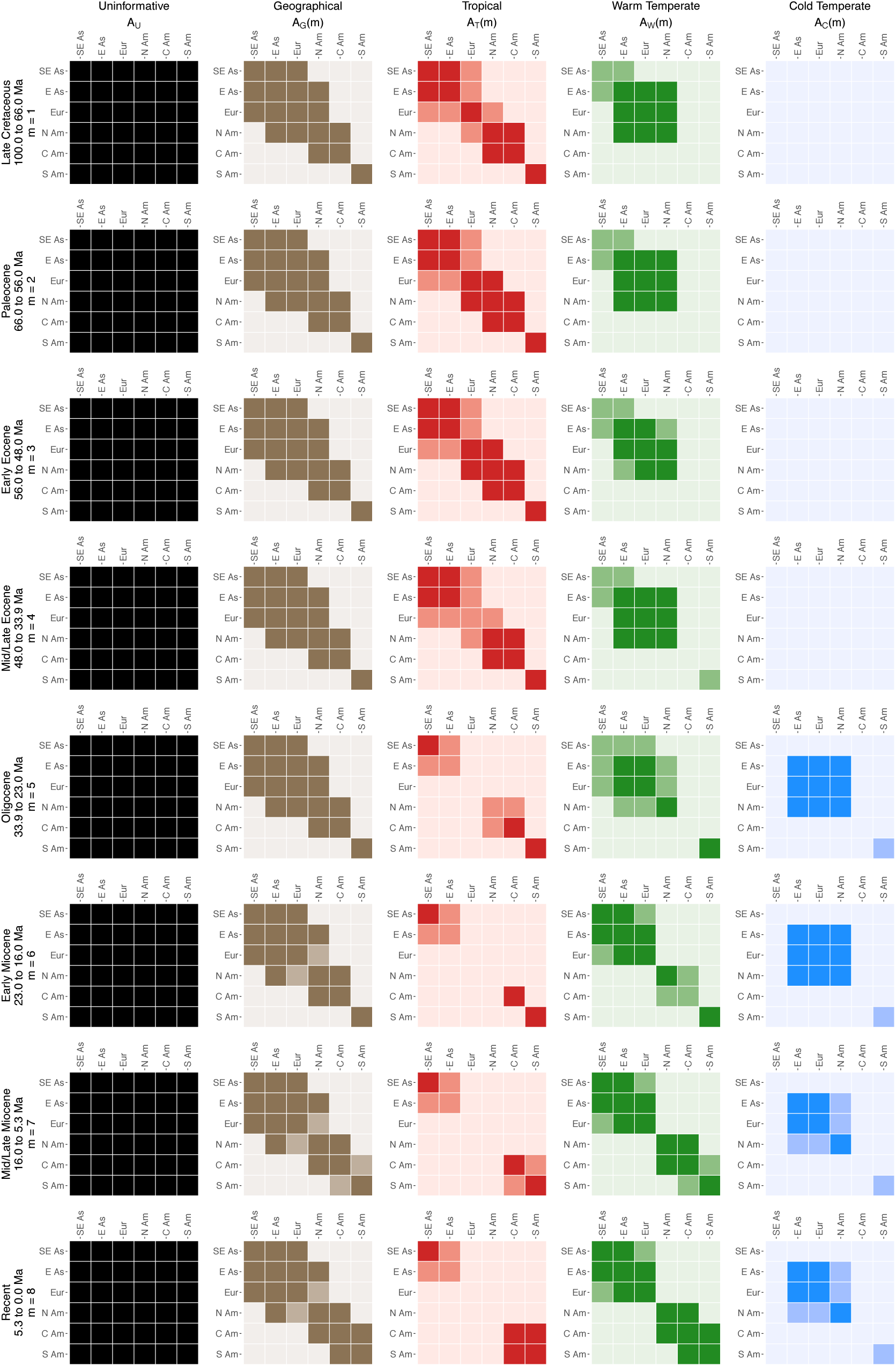
Availability and connectivity of biomes from Late Cretaceous (100 Ma) to Recent. Adjacency matrices are used to structure the time-stratified phylogenetic biome shift process. Rows correspond to eight major time intervals, while columns correspond to regional features, specifically uninformative features (black), simple geographical features (brown), or features involving the tropical (red), warm temperate (green), and cold temperate (blue) forest biomes. The matrix for each time and feature encodes the availability of (the diagonal) and the connectivity between (off-diagonal) regions for that feature at that time, where matrix rows and columns correspond to source and destination regions, respectively. Cells representing availability and connectivity are shaded to represent those features as being strong (dark), weak (medium), or marginal (light).

The BIOME4 reconstructions we obtained are 1-degree global rasters that classify each cell (location) as one of 27 possible biomes. BIOME4 biomes are inferred from assorted botanical and climatological information (observed or imputed) that represents a given time interval. For each BIOME4 reconstruction, we interpreted the 11 forest biomes as approximations of the availability and connectivity of mesic forest biomes among regions, then compared those features to our literature-based reconstructions. Previously published BIOME4 reconstructions were available for times corresponding to the Early-Mid Eocene (Herold et al. 2014), the Late Eocene and the Oligocene (Pound and Salzmann 2017), the Mid/Late Miocene (Pound et al. 2011, 2012), and the Pliocene (Salzmann et al. 2008; Salzmann, Haywood, and Lunt 2009). For time intervals that lacked published BIOME4 reconstructions, we compared our paleobiome maps to unpublished reconstructions provided by P. J. Valdes (pers. comm.) that were built from proprietary data. Our impression was that BIOME4 reconstructions tended to infer arid biomes that were more expansive and forest biomes that were less expansive than was generally supported by the paleobotanical literature. For example, the BIOME4 reconstruction inferred that the Amazon contained more xeric shrubland than tropical forest at the present. As such, we took presences of mesic forests under BIOME4 as conservatively restricted estimates of their true distributions. That being said, the majority of literature-based and BIOME4 reconstructions shared two important features: (1) the high degree of availability and connectivity among tropical and warm temperate biomes in the Northern Hemisphere before the Oligocene, and (2) the sudden rise of cold temperate forests concomitant with the disappearance of many tropical forests, beginning with the Oligocene.

We classified the availability and connectivity of biomes within regions into three categories of features—strong, weak, and marginal—that were appropriate to the scale of the regions and the precision of the ancestral biome estimates. Biomes with a strong presence displayed ≥25% regional coverage, biomes with a weak presence covered <25% of a region, while biomes with marginal presence covered <1% of a region. Likewise, the connectivity of a biome between two regions at a given time is scored as either strong, weak, or marginal, depending on how continuously biomes are inferred to have been distributed near regional adjacencies. For example, the modern distribution of the cold temperate biome throughout East Asia and Europe is consistent with strong connectivity, while North America is only weakly connected to cold temperate Eurasian biomes through its fragmented and transient arctic connections. Independent of the distribution of biomes, we similarly scored the geographical connectivity between regions as strong, weak, and marginal, using the equivalent of the modern connection between Central and South America through the Isthmus as Panama to minimally qualify as strong connectivity, and distances between modern Europe and North America to represent weak connectivity. The geographical connectivity between two regions is, generally, at least as strong as the strongest biome-dependent connection for those same regions. Together, the availability and connectivity for each region, each biome, and each time interval is encoded into a series of paleobiome graphs, which we later use to define the rates at which biome shift and dispersal events occur.

Our paleobiome graphs capture several important aspects of how mesic forest biomes moved and evolved (Fig. 2). The Late Cretaceous through the Paleocene and Early Eocene was a prolonged period of warm, wet conditions during which the poles had little to no ice. Throughout that time, tropical forests were prevalent in all six of our regions, while warm temperate forests were widespread only throughout East Asia, Europe, and North America. The Eocene witnessed the emergence of a so-called boreotropical flora, which contained a curious mixture (from our modern vantage point) of tropical and temperate plant genera (Wolfe 1975; Tiffney 1985a, b). This largely broadleaved evergreen forest type appears to have spread widely around the northern hemisphere via the Beringia Land Bridge, the North Atlantic Land Bridge, and the Tethys Sea (Tiffney 1985 a, b; Wolfe 1985; Morley 2000; Willis and McElwain 2014; Graham 2011, 2018; Baskin and Baskin 2016), and to have persisted through the Mid/Late Eocene. With the Oligocene, the opening of the Drake Passage and the closure of the Tethys Sea redirected global ocean currents. Together with steep declines in atmospheric CO_2_ levels, this ushered in cooler and drier conditions worldwide. The ensuing global climatic changes progressively restricted tropical forests to more equatorial regions, inducing the disjunction we find among modern tropical forests (Latham and Ricklefs 1993; Wiens and Donoghue 2004; Donoghue 2008). As the boreotropical forests receded, they were first replaced by warm temperate forests, and then eventually by cold temperate and boreal forests. Following this global revolution of biome structure, connectivity between Old World and New World tropical forests never again matched that of the Paleocene-Eocene boreotropical beltway.

We illustrate how a lineage might evolve with respect to different distributions of biomes within and between regions over time with the aid of Figure 2. A lineage that freely disperses between regions and shifts between biomes, regardless of the historical condition of the planet, might transition between regions under fully connected matrices (Uninformative, first column). Lineages that are only dispersal-limited by terrestrial connectivity disperse under the adjacency constraints encoded in the second column of matrices (Geographical, second column). However, lineages that are dispersal-limited by biome availability and connectivity might disperse according to the paleobiome patterns shown in the third, fourth and fifth columns (tropical, T; warm temperate, W; and cold temperate, C). For example, a lineage that is strictly adapted to the warm temperate biome would disperse according to the warm temperate series of paleobiome graphs (fourth column). If that lineage shifted its affinity from a warm temperate to a tropical biome, that lineage would thereafter shift between biomes and disperse between regions under the adjacency matrix structures of the tropical biome (third column) until the lineage next shifted biomes. However, biome shift rates also should depend on what biomes are locally accessible. For example, a North American lineage would have the geographical opportunity to shift from warm temperate into tropical biomes during the Paleocene, an epoch when both biomes have strong presences in North America. But North American tropical forests decline and then disappear throughout the Oligocene and Miocene, extinguishing the opportunity for such a biome shift during more recent times. The next section formalizes how we model the complex interactions between biomes, regions, phylogeny, and time with these dynamics in mind.

### A time-stratified regional biome shift model

The regional biome shift process may be viewed as a model that defines the interactions (if any) of its two subprocesses, the biome shift process and the dispersal process. We model biome shifts using a simple continuous-time Markov chain (CTMC) with time-stratified rates (i.e. piecewise constant time-heterogeneous rate matrices; Ree et al. 2005; Buerki et al. 2011; Bielejec et al. 2014; Landis 2017). Because transition rates between regions depend in part on a lineage’s biome affinity, and rates of shifting between biomes depend in part on a lineage’s geographical location, the two characters do not evolve independently. To impose interdependence between biomes and regions, we define a rate matrix over the compound state space by modifying the approach of Pagel (1994), while also drawing on insights pioneered in newer trait-dependent models of discrete biogeography (Sukumaran et al. 2015; Sukumaran and Knowles 2018; Matos-Maravı et al. 2018; Lu et al. 2019; Klaus and Matzke 2019).

Accordingly, we define the CTMC to operate on the compound biome-region state, (*i*, *k*), where *i* is the biome and *k* is the region. With this in mind, our goal is to compute the probability of a lineage transitioning from biome *i* in region *k* to biome *j* in region *l*, or (*i*, *k*) into (*j*, *l*). First, we take *β*_i,j_ to model the shift rate between biomes *i* and *j*, and *δ*_k,l_ to model the dispersal rate between regions, *δ*_k,l_. Importantly, the values of *β* and *δ* themselves do not directly depend on time. We eventually multiply these base rates by time-dependent paleogeographical and paleoecological factors represented in our time-stratified (or epoch) model.

Computing the transition probabilities for a time-stratified model requires that we define an instantaneous rate matrix *Q*(*m*) for any supported time interval, *m*. Following Landis (2017), we define the rate matrix *Q*(*m*) as the weighted average of several rate matrices, each capturing different paleogeographical features

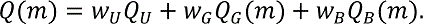

The three matrices on the right-hand side of Equation 1 are the (paleogeographically) uninformative rate matrix, *Q*_U_, the geographical rate matrix, *Q*_G_, and the biome rate matrix, *Q*_B_. In reference to Figure 2, we wish to learn the relative influence of the uninformative (first column), geography (second), and biome (third, fourth, or fifth) matrix features on the biome shift process. Supplement 1 demonstrates how rate matrix values are computed for a toy example with two regions and two biomes.

The first rate matrix (*Q*_U_) may be considered as an uninformative rate matrix that sets the relative transition rates between all pairs of regions, and separately between all pairs of biomes, as time-independent and context-independent,

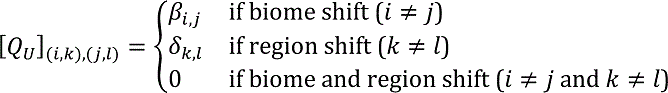

The effect is that biome shifts between biomes *i* and *j* follow the rates *β*_i,j_ and dispersal events follow the rates *δ*_k,l_ regardless of the age of a lineage or the lineage’s biome-region state. As we develop rate matrices for geography (*Q*_G_) and biomes (*Q*_B_) below, the second role for *Q*_U_ is that it allows for lineages to disperse or shift regardless of whether the connectivity/availability of the involved regions and/or biomes are scored as strong, weak, or marginal.

Because we do not know precisely what, if any, influence strong, weak, and marginal features should exert upon the dispersal or biome shift processes, we estimate the relative effects of those features across all adjacency matrices, the geographical adjacency matrix,

*A*_G_, and the biome adjacency matrices, *A*_E_, *A*_F_, and *A*_G_, for the tropical, warm temperate, and cold temperate biomes, respectively (Fig. 2). Specifically, we assign the fixed values of 0 and 1 to marginal and strong features, respectively, then assign all weak features one shared intermediate value, estimated as 0 < y < 1. In effect, y controls the degree of contrast of medium-colored cells across all adjacency matrices in Figure 2. For example, tropical connectivity during the Late Cretaceous is represented by the adjacency matrix

*A*_E_(1). At that time, tropical connectivity between East and Southeast Asia is strong

([*A*_E_(1)]_IJK,LIJK_ = 1), connectivity between East Asia and Europe is weak

([*A*_E_(1)]_IJK,INO_ = y), and connectivity between East Asia and South America is marginal

([*A*_E_(1)]_IJK,LJP_ = 0). Future studies may find that one can model and reliably estimate separate values of y for different features, even though we do not investigate this possibility at present.

The second rate matrix (indexed G for geography, *Q*_G_) is structured according to biome-independent paleogeographical features, such as the simple terrestrial connectivity between regions. Connectivity is encoded as either as strong, weak or marginal in the adjacency matrix, *A*_<_(*m*).

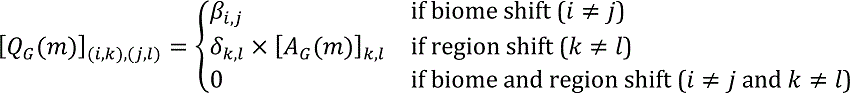

The third rate matrix (indexed B for biome, *Q*_B_) defines the shift rates between biomes and the dispersal rates between regions to depend on the spatiotemporal distribution of biomes. A lineage’s biome shift rate depends on whether the receiving biome, *j*, has a strong, weak, or marginal presence in the region it currently occupies, *k*. Likewise, the dispersal rate for a lineage that is currently adapted to biome type *i* depends on whether the source region, *k*, and destination region, *l*, share a strong, weak, or marginal connection.

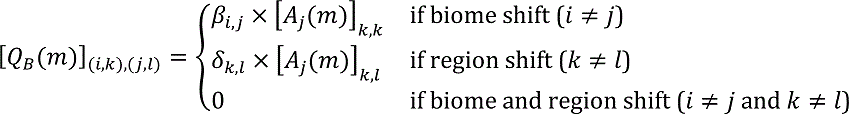

It is crucial to recognize that *Q*_B_(*m*) defines shift rates involving biome *j* to depend on the adjacency matrix for biome *j* during time interval *m*. This key property means that lineages currently adapted to biome *j* disperse with rates according to the interregional connectivity of biome *j*, and lineages newly adapting to biome *j* do so at a rate depending on the local availability of biome *j*.

The transition rates (and probabilities) between biome-region pairs are not expected to be symmetric or equal across time intervals. For example, if biome *j* first appears in region *k* during time interval *m* + 1, then we would potentially see an increase in the biome shift rate into biome *j*, i.e. [*Q*(*m*)]_(2,5),(3,5)_ < [*Q*(*m* + 1)]_(2,5),(3,5)_. Nor are transition rates necessarily symmetric or equal within a given time interval. If region *k* contains biome *i* during time interval *m*, but region *l* does not, then we might find that lineages adapted to biome *i* disperse less easily from *k* into *l* than *l* into *k*, i.e.

[*Q*(*m*)]_(2,5),(2,6)_ < [*Q*(*m*)]_(2,6),(2,5)_. Similarly, if region *k* contains biome *i* but not biome *j* during time interval *m*, then lineages inhabiting region *k* might tend to shift less easily from biome *i* into *j* than from *j* into *i*, i.e. [*Q*(*m*)]_(2,5),(3,5)_ < [*Q*(*m*)]_(3,5),(2,5)_.

Fluctuating asymmetries in the rates over time means that each biome-region state may exhibit different source-sink dynamics across that timescale. During a period of low accessibility, a biome-region state might rebuff immigrants and lose occupants when acting as a source, but then gain and retain inhabitants during a later phase should that biome-region become a local refugium when acting as a sink (Goldberg et al. 2005). These fluctuating source-sink dynamics may be characterized by the stationary distribution, which defines the expected proportion of lineages found in each biome-region state assuming lineages evolve along an infinitely long branch within a given time interval.

Biome-regions that are easy to enter and difficult to leave tend towards higher stationary probabilities for a given time interval.

We approximate the stationary probability for biome *i* in region *k* during time interval *m* with

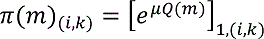

where *μ* is a rate taken to be sufficiently large that the evolutionary process reaches stationarity. Note that the bracketed term on the righthand side of the equation is the transition probability matrix for changes between biome-region pairs. In theory, when *μ* is large, all rows in this matrix have arbitrarily similar transition probabilities, which lets us take any row (e.g. the first row) to represent the stationary probabilities.

The time-dependent source-sink dynamics in Figure 3 show how the availability of and connectivity between regional biomes structure each time interval’s stationary distribution. Stationary probabilities before the Oligocene tend to favor tropical biomes in all regions but favor cold temperate biomes afterwards. This means that if the historical spatial structure of biomes is relevant to biogeography, then lineages originating in the Paleogene would more likely be adapted to tropical than to cold temperate forests simply because cold temperate forests were a more marginal biome during that period of Earth’s history.

**Figure 3.**
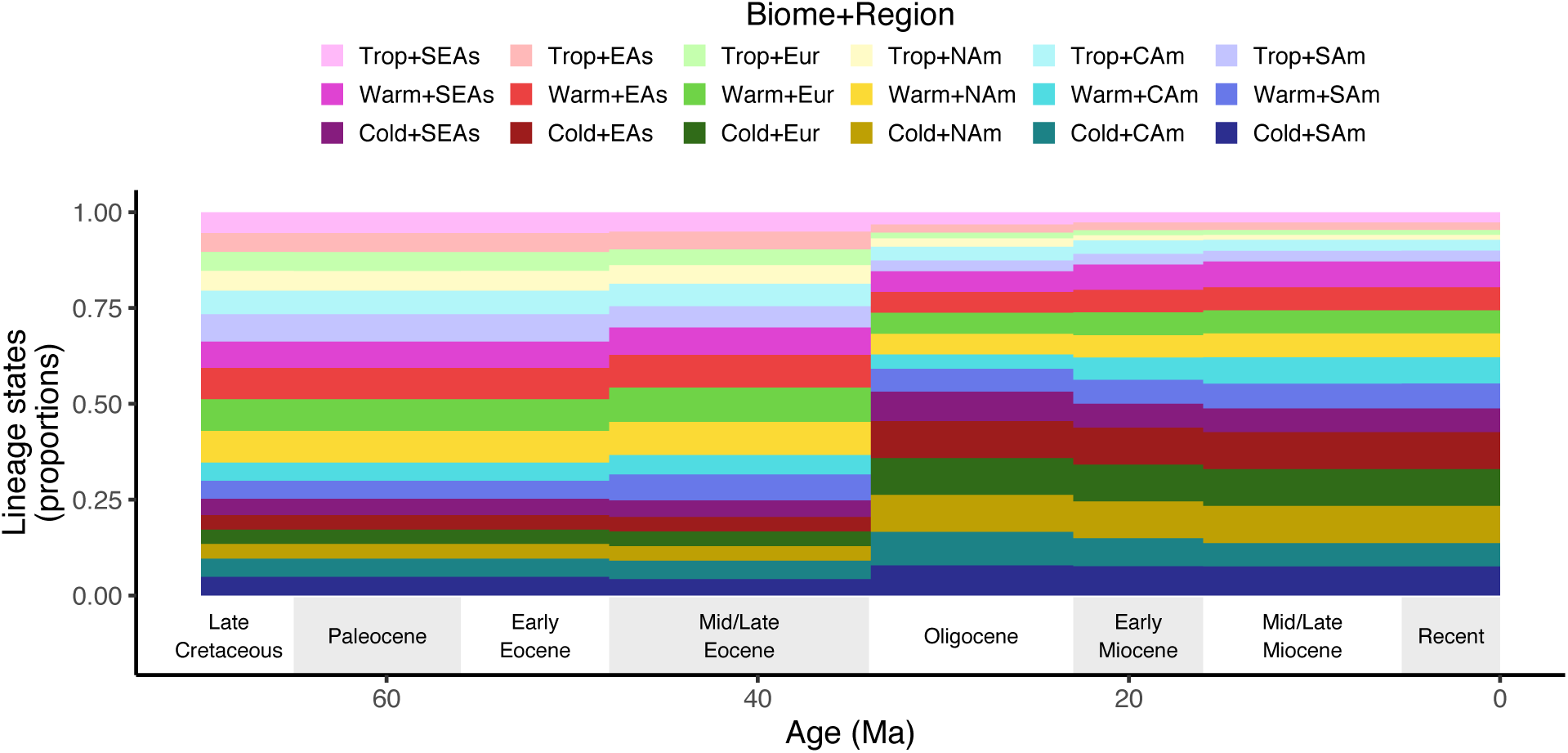
Stationary distribution of biome-region states under the paleobiome model. The stationary probabilities across biome-regions (y-axis) vary with respect to time (x-axis). Stationary probabilities were computed assuming that biome and region shifts are expected to occur in equal proportion (*β* = *δ* = 0.5), that lineages tend to shift and disperse in a manner than depends heavily on the paleobiome structure (*w*_U_ = 0.04, *w*_G_ = 0.16, and *w*_B_ = 0.8), and that biomes with strong presences primarily define the structure of biome graphs (*y* = 0.1). Parameters were chosen to show interesting variation. Note, all stationary probabilities would be equal over all times if *w*_U_ = 1.

We can now completely define the time stratified rate matrix, *Q*(*m*), and the stationary frequencies at the root of a phylogeny, *π*(*m*_root_), where *m*_root_ is the time interval corresponding to the root node age. Together, these model components let us compute the probabilities of lineages transitioning from one biome-region pair to another while accounting for the spatiotemporal dynamics of biomes, and thus compute the phylogenetic model likelihood with the discrete state pruning algorithm (Felsenstein 1981).

Now that we have fully defined the model, there are several implicit properties that are worth stating explicitly. First, a lineage cannot both shift its biome affinity and disperse into a new region in the same moment of time; one event is needed for each transition, and so event order matters. While we think this assumption is reasonable in many cases, including in *Viburnum*, it is certainly conceivable that region and biome shifts could occur simultaneously. We elaborate on this point in the Discussion, but note here that it would be possible to explore the effect of modeling such paired transitions by adapting parameterizations designed for multiple-nucleotide substitution codon models (Koisol et al. 2007). Second, the relative importance of the matrix feature weights (*w*_U_, *w*_G_, *w*_B_) and of the weak availability/connectivity weight (y) are estimated from the data: the matrix *Q*(*m*) reduces to the uninformative matrix, *Q*_U_, when *w*_U_ = 1, while the importance of the historical structure of biomes is most pronounced when *w*_B_ is large compared to other *w* and *y* parameters. Third, the process models lineages as being predominantly present in a single region and biome at a time without influencing speciation or global extinction rates, which are assumptions we made both to simplify the exposition of the method, but also to reduce computational burden. The Discussion brings more attention to all of these properties.

### Bayesian inference

The Bayesian posterior density was estimated using the Markov chain Monte Carlo (MCMC) algorithm implemented in RevBayes (Höhna et al. 2016). RevBayes scripts and datasets are available at http://github.com/mlandis/biome_shift. The first 10% of posterior samples were discarded as burn-in. All parameter estimates have effective sample sizes well over 200. Two independent chains were run per analysis to verify MCMC convergence. We analyzed our data under three model settings: the *Paleobiome* setting that used the time-heterogeneous graphical structure presented in Figure 2; the *Modern Biome* setting that used the graphical structure from the Recent time interval to represent all time intervals; and the *Null Biome* setting that ignored regional and biome structure by fixing *w*_U_ = 1.

Departing from the general model description above, we reparameterized our applied model to eliminate informative priors wherever possible. This helped ensure that our posterior estimates are driven by the data through the likelihood function, not the prior (discussed further in Supplement 2). We assigned uninformative prior distributions to our graph weights, (*w*_U_, *w*_L_, *w*_B_) ∼ Dirichlet(1, 1, 1), and to our weak feature strength parameter, y ∼ Uniform(0, 1). We treated each biome shift rate as an independently estimated parameter, *β*_2,3_ ∼ Uniform(0, 1), although we fixed the biome shift rates between tropical and cold temperate biomes equal to zero. Fixing those transition rates to zero both reduces the number of free parameters and reflects the observation that, analogous to the geographical structure of regions, biomes have an important ecological and climatological structure (Whittaker 1970). In *Viburnum*, this assumption is reasonable, as the clade contains no tropical-cold temperate sister species pairs, and previous ancestral biome estimates for the group did not confidently infer any shifts between those biomes (Landis et al. 2020). Because we constrained biome-independent dispersal between regions through graphical structures (*Q*_G_) and weight parameters (*w*_U_and *w*_G_), we fixed the relative dispersal rate to *δ*_k,l_ = 1 (which is potentially rescaled by *Q*_G_ and *w*_G_). Thus, the relative biome shift rates *β* and dispersal rates *δ* all have values between 0 and 1. To model the relative proportion of biome shifts to dispersal events, we multiply *β* by the factor _f__β_ ∼ Uniform(0,1) and multiply *δ* by _f__δ_ = (1 − *f*_β_). Finally, we rescaled the instantaneous rate matrix, *Q*, for the entire evolutionary process by a global clock parameter, *μ* ∼ LogUniform(10^ef^, 10^Y^), that is uniformly distributed over orders of magnitude. For the *Viburnum* analysis, we used Bayes factors (Jeffreys 1935) to compare the relative fit of the *Paleobiome*, *Modern Biome*, and *Null Biome* models to the data. Power-posterior distributions were approximated with a parallelized MCMC sampler (Höhna et al. 2017).

Marginal likelihoods were estimated from those power-posterior samples using the stepping stone algorithm (Xie et al. 2011). Our analysis produced several inferences, which we summarized in various ways.

Ancestral states and stochastic mappings (Huelsenbeck et al. 2003) were generated during MCMC sampling by first drawing ancestral state estimates for all nodes in phylogeny, then sampling evolutionary histories using a uniformization method (Rodrigue et al. 2008) that was adapted for time-stratified models (Landis et al. 2017). Ancestral state estimates show the posterior probabilities for biome-region states for each node. Lineage-state proportions through time were computed from the posterior distributions of stochastically mapped histories. We computed the posterior mean count of lineage-states through time as the number of lineages in each state for each time bin across posterior samples divided by the total number of posterior samples. Lineage-state counts were converted into lineage-state proportions by dividing each count by the total number of lineages in that time bin to give proportions that lie between 0 and 1. In addition, we classified whether or not each lineage-state for each time bin was congruent with any locally prominent biome as defined by the paleobiome graph (Fig. 2). Each binned state was labeled as a *biome mismatch* if the lineage’s biome had only a marginal presence in the lineage’s region. Otherwise, the state was labelled as a *biome match*. To summarize these results, we also computed the proportion of tree length where lineage states match or mismatch paleobiome structure in three ways: for the total tree length, for tree length before the Oligocene (>34 Ma), and for tree length after the Oligocene (≤ 34 Ma). To assess whether the use of the *Paleobiome* model unduly biased posterior estimates (e.g. by inflating the prior expectations of biome match proportions), we generated stochastic mappings conditional on the phylogeny and tip states while sampling model parameters from the prior (details given in Supplement 2).

We were also interested in the ordered *event series* that resulted in major transitions between biomes and regions. For biomes *A*, *B*, and *C* and regions *X*, *Y*, and *Z*, we named the six series patterns for pairs of events. Series in which species shift biomes and then disperse (*AX* → *BX* → *BY*) are called *biome-first* event series. In contrast, *region-first* series have dispersal followed by a biome shift event (*AX* → *AY* → *BY*). The remaining four event series involve two consecutive biome shift or two dispersal events. *Biome reversal* (*AX* → *BX* → *AX*) and *region reversal* (*AX* → *AY* → *AX*) sequences indicate event series in which the lineage departs from and then returns to its initial state (*AX*). Analogously, *biome flight* (*AX* → *BX* → *CX*) and *region flight* (*AX* → *AY* → *AZ*) sequences are recognized by series of two biome shifts or two dispersal events that leave the lineage in a new state (*CX* or *AZ*) relative to the lineage’s initial state (*AX*). We computed the proportion of each series type for a single posterior sample by classifying stochastically mapped state triplets (event series of length two) in our phylogenetic tree using a simple root-to-tip recursion.

To start the recursion, we took the stochastically mapped root state to be the second state in the triplet, *X*_Ooop_, then sampling the preceding state, *X*_KNqOoop_, from the sampling distribution obtained by Bayes rule

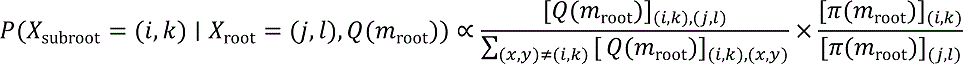

where *Q*(*m*_root_) is the root node’s rate matrix and *π*(*m*_root_) is its stationary distribution with values determined by the evaluated posterior sample. Following that, we traversed towards the tips of the tree to collect changes in the stochastic mapping for each lineage’s biome-region state, classifying the state triplet’s type, and updating the triplet states appropriately (i.e. the new second and third states replace the old first and second states) with each step of the recursion.

Finally, we wished to examine if and how the distribution of evolutionary events and event series changed with time under alternative assumptions about the biome structure. We were particularly interested in two classes of event proportions: proportions of various types of biome shift and dispersal events, and proportions of the various types of event series. To estimate the posterior proportions of biome shift and dispersal event types through time, for each posterior sample, we divided the count for each distinct biome shift and dispersal event type by the total number of events within each major time interval.

Event proportions from the Late Cretaceous could not be reliably estimated, as the two *Viburnum* lineages that existed between the origin of the group (∼70 Ma) and the end of the Late Cretaceous (∼66 Ma) rarely changed in biome-region state. Although we normalized our event proportions using all 126 distinct dispersal and biome shift event types, our results only display the four biome shift and four dispersal event types among all combinations of the warm and cold temperate forests of East Asia and North America. In a similar manner, we computed the posterior proportions for all six types of event series, using the time of the second event in each series for each series age.

We provide the posterior means and the 80% and 95% highest posterior densities (HPD80 and HPD95, respectively) for each time interval, event or event series type, and biome structure model. Constructing Bayesian credible intervals for these proportions is somewhat unusual, since the proportions are ratios of counts (i.e. not all non-negative real numbers are supported). As a result, several HPDs share the exact same bounds, particularly when the total number of events is low (e.g. values such as 1/2 or 1/3).

### Simulation experiment

We measured how reliably we can select models in which biome structure influences the biome shift process (*w*_=_ > 0) for *Viburnum* with simulated data. All simulations assumed the same *Viburnum* phylogeny used in the empirical example and used the same biome and regions designated by the paleobiome structure model. We simulated data under five conditions that primarily adjusted the relative weight for *w*_B_, named: null effect, where (*w*_U_, *w*_G_, *w*_B_) = (1,0,0); weak effect, where (*w*_U_, *w*_G_, *w*_B_) = (1,2,4)/7; medium effect, where (*w*_U_, *w*_G_, *w*_B_) = (1,2,8)/11; strong effect, where (*w*_U_, *w*_G_, *w*_B_) = (1,2,16)/19; and very strong effect, where (*w*_U_, *w*_G_, *w*_B_) = (1,2,32)/35; with each denominator ensuring the weights sum to 1. For all conditions, we assumed _f__b_ = 0.75, _f__δ_ = 0.25, and *y* = 0.1. Biome shift rates were set to equal 1, except transitions between cold temperate and tropical forests, which were set to 0. The event clock was set to *μ* = 0.03, except for the null condition, which was assigned a slower rate of *μ* = 0.01 to account for the fact that fewer event rate penalties are applied to it than the non-null conditions.

We then simulated 100 replicate datasets in RevBayes for each of the four conditions under the regional biome-shift model described above, and estimated the posterior density for each simulated dataset using MCMC in RevBayes.

We were primarily concerned with how our posterior estimates of *w*_B_ respond to differing simulated values for *w*_B_. To summarize this, we first report the posterior median values of *w*_B_ across replicates so they may be compared to the true simulating value. Next, we computed what proportion of our replicates select a complex model allowing *w*_B_ > 0 in favor of a simpler model where *w*_B_ = 0 using Bayes factors. Bayes factors were computed using the Savage-Dickey ratio (Verdinelli and Wasserman 1995), defined as the ratio of the prior probability divided by the posterior probability, evaluated at the point where the complex model collapses into the simpler model (i.e. *w*_B_ = 0, in our case). We approximated the posterior probability at this collapse-point by smoothing our posterior samples for *w*_B_ with a beta-kernel density estimator (Moss and Tveten, 2019). We interpret the strength of significance for Bayes factors as proposed by Jeffreys (1961), requiring at least ‘Substantial’ support (BF > 3) to select the more complex model (*w*_B_ > 0).

## RESULTS

### Simulation experiment

Simulated datasets yielded larger estimates of *w*_B_ and more soundly rejected null models (*w*_B_ = 0) as the effect strength *w*_B_ increased from Weak to Very Strong (Fig. 4). No datasets simulated under the Null condition (*w*_B_ = 0) signalled Substantial support (or greater) for the paleobiome-dependent model (*w*_B_ > 0), indicating a low false positive rate. Only 9% of datasets simulated under Weak effects (*w*_B_ = 4/7 ≈ 0.57) generated No support for the *w*_B_ > 0 model, while ∼32% of those replicates qualified as Substantial support or greater. Data simulated under the Moderate condition (*w*_B_ = 8/11 ≈ 0.73) rejected the simple model 57% of the time with at least Substantial support. Under Strong (*w*_B_ = 16/19 ≈ 0.84) simulation conditions, we selected models where *w*_B_ > 0 in 81% of cases, with Strong support in 65% of cases. Data simulated under Very Strong effects (*w*_B_ = 32/35 ≈ 0.91) generated support for models with *w*_B_ > 0 88% of the time, with over half of all replicates (54%) drawing Very Strong or Decisive support. Coverage frequency among simulations was consistently high across conditions, but with fairly wide credible intervals corresponding to the highest 95% posterior density (Fig. 4A). Because the posterior probability of *w*_B_ = 0 is used to approximate Bayes factor ratios, their relationship is made apparent by noting that the density of HPD95 lower bound estimates close to the value *w*_B_ = 0 (Fig. 4A) is correlated with the proportion of simulations that award no support to the *w*_B_ > 0 model (Fig. 4B).

**Figure 4.**
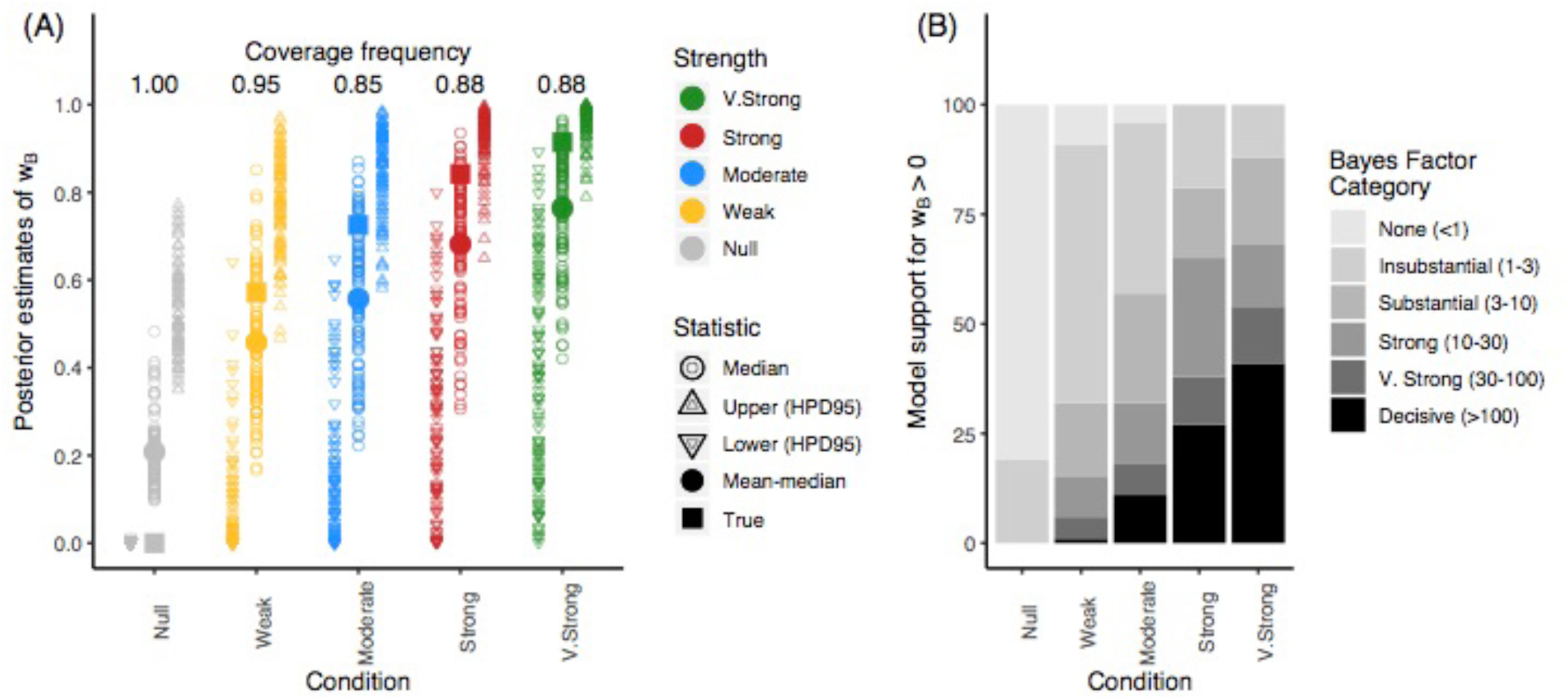
Simulation experiment results. One hundred datasets were simulated under five conditions that varied the strength of *w*_B_, then fitted to the paleobiome model to assess model performance. (A) Markers show the true simulated strength for *w*_B_ (closed square), the posterior median values estimated from simulated replicates (open circles), the median of those posterior medians (closed circle), and the upper and lower bounds of the 95% highest posterior density (open triangles). The coverage frequency reports the proportion of simulation analyses in which the simulating value of *w*_B_ is falls within the 95% highest posterior density. (B) Bars report the proportions of simulated datasets that supported the model where *w*_B_ > 0, categorized by the strength of that support in terms of Bayes factors (Jeffreys 1961).

### Ancestral biomes for Viburnum

Although *Viburnum* likely originated in East Asia regardless of the biome structure model (*p* > 0.99), no model reconstructed a single ancestral biome affinity with probability greater than *p* > 0.95 (Fig. 5). Where the *Paleobiome* analysis inferred East Asian biome affinities that favored a warm temperate (*p* = 0.88) or tropical (*p* = 0.09) but not a cold temperate (*p* = 0.03) origin, the *Modern Biome* analysis favored a cold temperate (*p* = 0.67) then warm temperate (*p* = 0.31) origin for *Viburnum* while assigning negligible probability to a tropical origin (*p* = 0.01). Relative to the *Paleobiome* estimates, the *Null Biome* analysis also assigned higher probabilities towards colder biomes (warm, *p* = 0.52; cold, *p* = 0.45; tropical, *p* = 0.02). Early diverging *Viburnum* lineages tended to follow warm/tropical biome affinities under the *Paleobiome* analysis or the cold/warm affinities under the *Modern*/*Null Biome* analyses before the Oligocene (>34 Ma). During the Oligocene (34–22 Ma), when cold temperate forests first expanded, many nodes still retained the warmer (*Paleobiome*) or colder (*Modern Biome*) affinities characteristic of the corresponding biome structure model, such as the most recent common ancestor (MRCA) of *V. reticulatum* and *V. ellipticum* or the MRCA of *V. rufidulum* and *V. cassinoides*. Otherwise, most ancestral biome inferences were consistent across the three models, beginning with the Mid/Late Miocene (<16 Ma).

**Figure 5.**
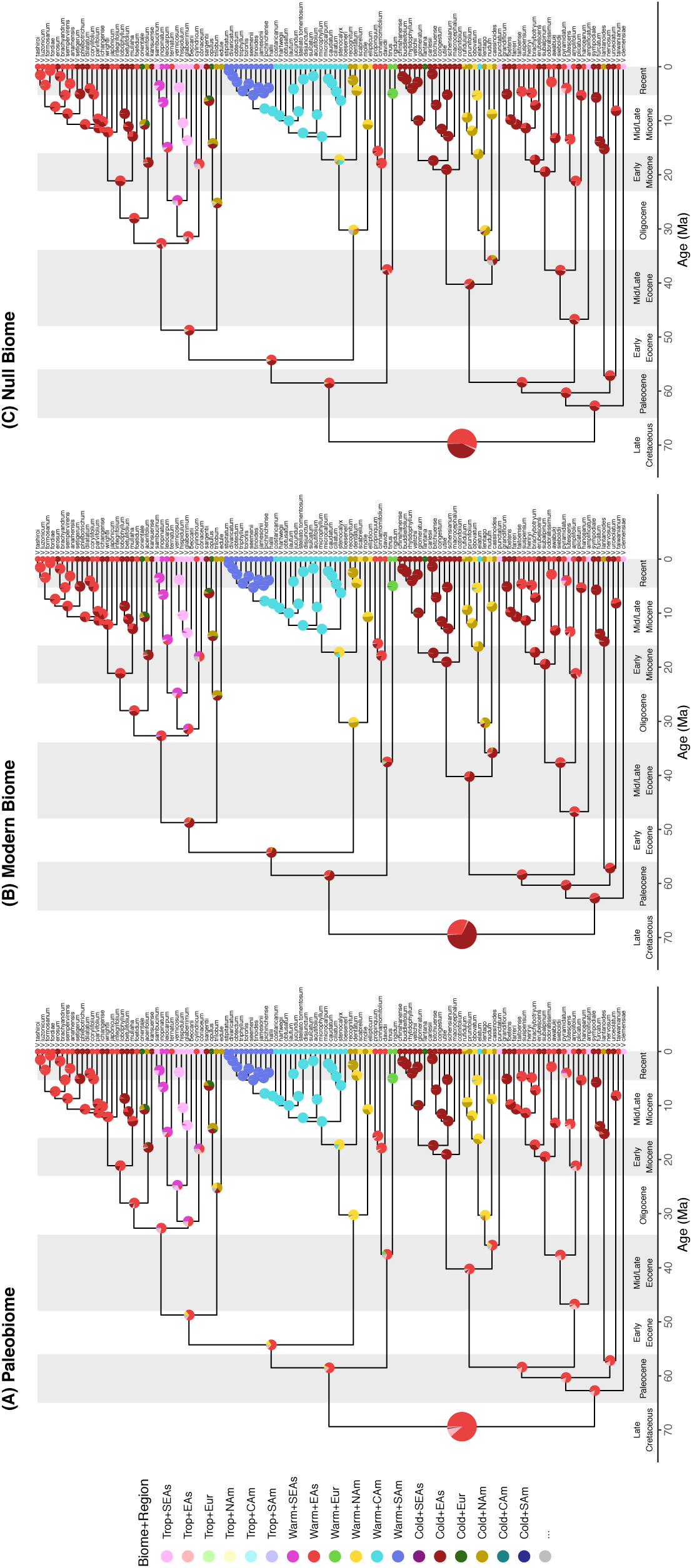
Ancestral biome-region state estimates for *Viburnum*. Estimates produced under (A) *Paleobiome*, (B) *Modern Biome*, and (C) *Null Biome* settings. Colored pie charts report posterior support for the three most probable biome-region states per node. Pie charts for root state probabilities are magnified to improve visibility. Vertical white and gray bands correspond to major geological timeframes referenced in this study.

The three biome structure models recovered different proportions of ancestral lineage-states through time, particularly before the Mid/Late Miocene (>16 Ma; Fig. 6A–C). Between the Paleocene and the Early Miocene, tropical lineages in East Asia and Southeast Asia constituted >20% diversity, declining to ∼12% of modern diversity under the *Paleobiome* analysis. Cold temperate lineages were nearly absent until the end of the Oligocene (34 Ma), but steadily rose to constitute roughly 25% of diversity by the Early/Mid Miocene (ca. 20 Ma). By comparison, *Modern Biome* estimates enriched the proportion of cold temperate viburnums, while reducing support for warm temperate and nearly eliminating support for tropical origins; tropical lineages remained in comparatively low proportion until the Miocene (< 22 Ma). The *Null Biome* analysis estimated proportions of warm and cold temperate lineages similar to those of the *Modern Biome* analysis from the Late Cretaceous (100 Ma) until the Oligocene (34 Ma), but with more Southeast Asian warm temperate lineages throughout.

**Figure 6.**
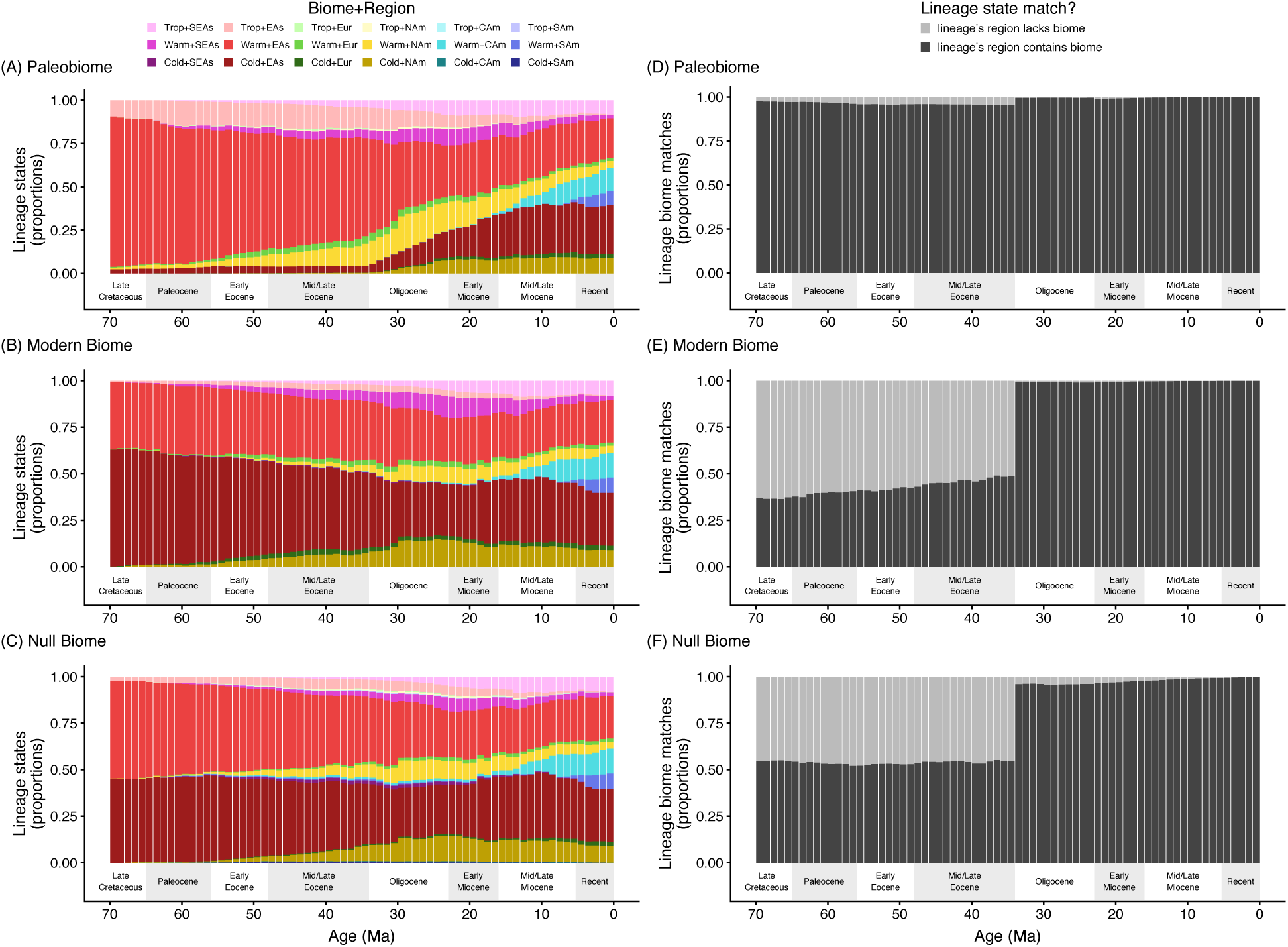
Ancestral proportions of lineage state frequencies through time for *Viburnum*. The left column (A–C) shows the lineages biome-region states, where regions differ by color and biomes differ by shading (see legend). Proportions of reconstructed lineages in each biome-region state are shown for estimates under the *Paleobiome* (A), *Modern Biome* (B), and *Null Biome* (C) settings. The right column (D–F) shows the proportion of lineages with biome states that match (dark) or mismatch (light) the non-marginal biomes that are locally accessible given any lineage’s location, as defined under the Paleobiome structure (see main text for details). Proportions of reconstructed lineages with biome match and mismatch scores are shown for estimates under the *Paleobiome* (D), *Modern Biome* (E), and *Null Biome* (F) settings.

For what proportion of time did lineages have biome affinities that were congruent with locally accessible biomes? Biomes rarely mismatched between lineages and regions under the *Paleobiome* setting (1.1% of tree length), with the mismatches increasing under the *Modern Biome* (8.6%) and *Null Biome* (8.7%) settings. Lineages were most often mismatched with their regions’ biomes before the Oligocene (Figs. 6D–F), where the pre-Oligocene proportion of mismatched branch lengths was always higher (*Paleobiome*, 5.8%; *Modern Biome*, 52.6%; *Null Biome*, 47.1%) than the post-Oligocene proportion (*Paleobiome*, 0.3%; *Modern Biome*, 0.8%; *Null Biome*, 1.7%) or the treewide proportions (above). It is unlikely that our posterior inferences are artifacts caused by undesirable interactions between our prior model and our biome structure models, since the prior-like distributions of stochastic mappings that we generated under the *Paleobiome*, *Modern Biome*, and *Null Biome* settings were nearly indistinguishable from one another (Supplement 2; Fig. S1).

To illuminate why the *Paleobiome* analysis produces distinctly warmer ancestral biome estimates, we turn to the fitted stationary probability for the root state, *π*(*m*_root_), (Fig. 7). Within East Asia, root node stationary probabilities estimated under the *Paleobiome* setting favored warm temperate or tropical forests over cold temperate forests (*π*_Trop+EAs_ = 0.06, *π*_Warm+EAs_ = 0.10, *π*_Cold+EAs_ = 0.02). The *Modern Biome* stationary probabilities instead favored cold or warm temperate forests over tropical forests (*π*_Trop+EAs_ = 0.03, *π*_Warm+EAs_ = 0.07, *π*_Cold+EAs_ = 0.08). Like the *Modern Biome* analysis, stationary probabilities under the *Null Biome* setting tended towards cold or warm temperate forests (*π*_Trop+EAs_ = 0.04, *π*_Warm+EAs_ = 0.06, *π*_Cold+EAs_ = 0.06), noting that the stationary probability per biome is uniform across regions by the design of the model. While the posterior root node stationary probabilities varied across biome-regions depending on what biome structure model was assumed (Fig. 7), all corresponding prior probabilities are approximately equal (Supplement 2; Fig. S2), suggesting that any differences in the posterior estimates are driven by the data through the likelihood function, and not forced through an induced prior.

**Figure 7.**
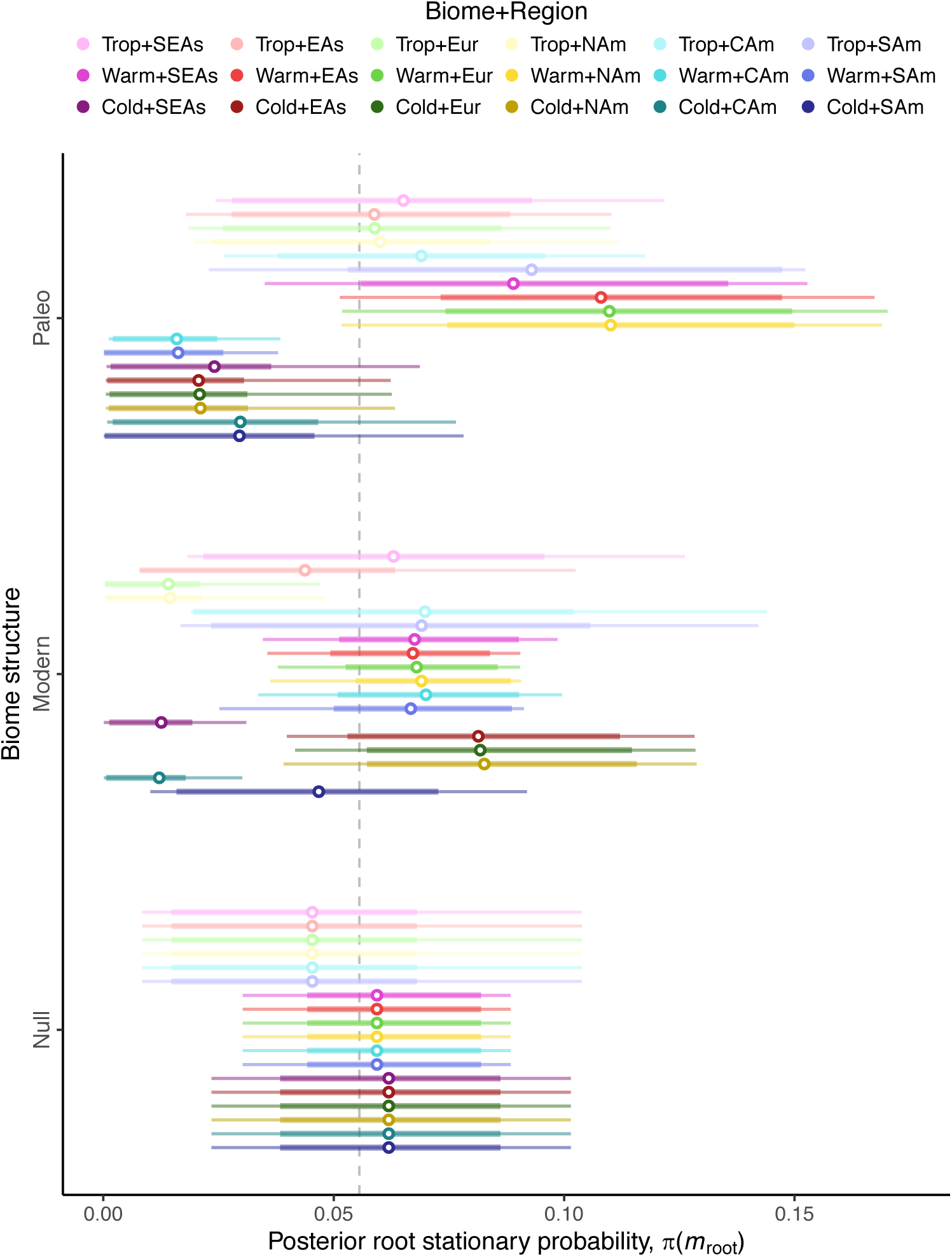
Stationary probabilities for the *Viburnum* root state during the Late Cretaceous. Posterior stationary probabilities for *π*(*m*_root_) are given for each biome structure model (grouped rows) and for each biome-region state (colors) as posterior means (points) and credible intervals (HPD80, thick lines; HPD95, thin lines).

Despite such differences between the *Paleobiome* and *Modern Biome* analyses in their ancestral state estimates and stationary probabilities, their parameter estimates for the base rate of change (*μ*), the proportion of biome shifts (_f__β_) to dispersal events (_f__δ_), and the graph weights (*w*_U_, *w*_G_, *w*_B_) were remarkably similar (Table 1). Both biome structure models estimate posterior means for *w*_B_ greater than 0.91; i.e., stronger in effect than assumed under the Very Strong simulation scenario (Fig. 4). Both models estimated credible intervals for *w*_B_ with lower bounds greater than 0.75 and posterior probabilities of *w*_B_ = 0 that were indistinguishable from zero, each corresponding to Decisive support for their respective biome structure models. More accurate Bayes factors estimated using marginal likelihoods unequivocally selected either the *Paleobiome* (BF_PN_ > 10^6^) and the *Modern Biome* (BF_MN_ > 10^6^) over the *Null Biome* structures for modeling biome shifts in *Viburnum*, but found only mild support in favor of the *Paleobiome* over the *Modern Biome* structure (BF_PM_ ≈ 1.8). Because inference under the *Null Biome* model set y = 1, posterior estimates of (*w*_U_, *w*_G_, *w*_B_) are indistinguishable from the prior. Parameter estimates for the relative biome shift rates differed across the three biome structure models, however. The *Paleobiome* estimates have relatively low cold-into-warm and high tropical-into-warm biome shift rates (posterior medians of *β*_EF_ = 0.67, *β*_FE_= 0.29, *β*_FG_= 0.81, *β*_GF_= 0.38) when compared to the *Modern Biome* (*β*_EF_= 0.50, *β*_FE_= 0.39, *β*_FG_= 0.81, *β*_GF_= 0.65) and the *Null Biome* estimates (*β*_EF_ = 0.54, *β*_FE_= 0.31, *β*_FG_= 0.74, *β*_GF_= 0.71).

**Table 1.** Regional biome shift parameter estimates. Posterior median estimates are in bold and 95% highest posterior densities are in brackets. Fixed parameters under the *Null Biome* analysis do not have brackets.

| Parameter | Biome structure |  |  |
| --- | --- | --- | --- |
|  | *Paleo* | *Modern* | *Null* |
| $\mu$ | **0.06**<br>[0.03,0.10] | **0.05**<br>[0.03,0.09] | **0.03**<br>[0.02,0.06] |
| $f_{\beta}$ | **0.85**<br>[0.75,0.94] | **0.83**<br>[0.69,0.93] | **0.92**<br>[0.85,0.97] |
| $f_{\delta}$ | **0.15**<br>[0.06,0.25] | **0.17**<br>[0.07,0.31] | **0.08**<br>[0.03,0.15] |
| $\beta_{TW}$ | **0.67**<br>[0.20,1.00] | **0.50**<br>[0.05,0.95] | **0.54**<br>[0.10,1.00] |
| $\beta_{WC}$ | **0.81**<br>[0.47,1.00] | **0.81**<br>[0.49,1.00] | **0.74**<br>[0.39,1.00] |
| $\beta_{WT}$ | **0.29**<br>[0.10,0.62] | **0.39**<br>[0.11,0.85] | **0.31**<br>[0.06,0.64] |
| $\beta_{CW}$ | **0.38**<br>[0.01,0.80] | **0.65**<br>[0.34,1.00] | **0.71**<br>[0.33,1.00] |
| $w_U$ | **0.01**<br>[0.00,0.07] | **0.02**<br>[0.00,0.08] | **1** |
| $w_G$ | **0.04**<br>[0.00,0.18] | **0.04**<br>[0.00,0.20] | **0** |
| $w_B$ | **0.94**<br>[0.78,1.00] | **0.93**<br>[0.76,1.00] | **0** |
| $y$ | **0.65**<br>[0.27,0.99] | **0.52**<br>[0.09,0.95] | **1** |

Finally, we found that the *Paleobiome* analysis estimated proportions of biome shift and dispersal events that are more temporally dynamic than those proportions estimated under the *Modern Biome* and *Null Biome* models (Fig. 8A–C). Under the *Paleobiome* estimates, dispersal events from East Asia into North America within the warm temperate biome were relatively common throughout the Late Eocene. With the onset of Oligocene cooling, biome shifts from warm into cold temperate forests in East Asia rose from low to high proportions to become the most frequent transition type. In contrast, event proportions under the *Modern Biome* and *Null Biome* analyses reconstructed high proportions of biome shifts between the warm and cold temperate forests of East Asia, ever since *Viburnum* first originated in the Late Cretaceous through the present. Paleocene dispersal of cold temperate lineages from East Asia into North America was also found to be relatively common when compared to the *Paleobiome* reconstruction. Regarding the event series proportions, biome reversal, biome-first, and region-first series were generally more common than biome flight, region flight, and region reversal series (Fig. 8D–F). The biome reversal event series was the most common event series type across all time intervals under the *Modern Biome* and *Null Biome* analyses, but not under the *Paleobiome* analysis. With the *Paleobiome* model, we found that the proportion of biome reversal series was lower, and the proportion of region-first series was higher, when compared to the other biome structure analyses, together creating a time interval between the Late Eocene and the Middle Miocene during which region-first events outpaced all other types of series.

**Figure 8.**
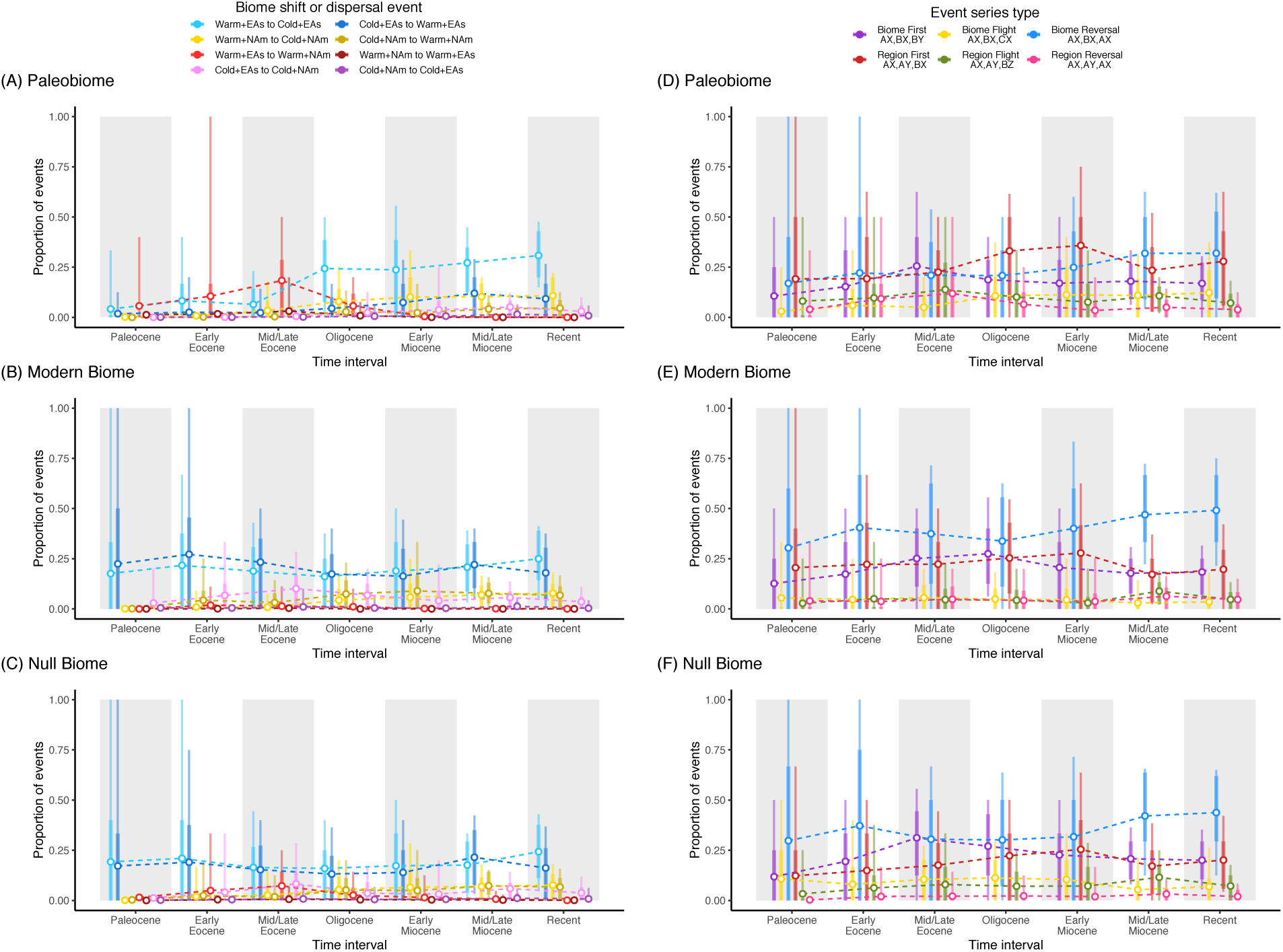
Proportions of inferred events and event series across major time intervals for *Viburnum*. Posterior proportions are presented as posterior means (points) and credible intervals (HPD80, thick lines; HPD95, thin lines). The left column (A–C) presents the proportions of estimated biome shift and dispersal events with respect to time, showing only the eight biome shift and dispersal events among the warm and cold temperate forests of East Asia and North America. Proportions of events are shown for inferences under the *Paleobiome* (A), *Modern Biome* (B), and *Null Biome* (C) settings. The right column (D–F) shows the proportions of the six types of event series with respect to time (defined in main text). Each event series type is labeled with a ‘state triplet’ to indicate either transitions in the biome (A, B, C) or region (X, Y, Z) state.

## DISCUSSION

The probability that a lineage will shift into a new biome is determined in part by geographical opportunities. Because the availability and connectivity of biomes varies across regions, evolutionary lineages do not share the same geographical opportunities to adapt to new biomes. Moreover, those geographical opportunities have changed as the spatial structure of Earth’s biomes evolved over time. As an evolutionary inference problem, the temporal dynamics of geographical opportunity is concerning: we typically infer ancestral biomes based on the phylogenetic distribution of biomes from extant species, yet their ancestors were likely exposed to geographical opportunities that were significantly (perhaps even radically) different from the opportunities of their living descendants.

Here, we have developed a Bayesian framework to model how phylogenetic lineages gain affinities with new biomes and disperse between regions in a manner reflecting the historical configuration of biomes through space and time. To do so, we modeled a time-stratified regional biome shift process using continuous-time Markov chains. The model is parameterized to allow biome shift and dispersal rates to depend on empirically structured paleobiome graphs, where each graph describes the availability and connectivity of biomes among regions within a given time stratum. We conducted a simple simulation experiment to show that we can identify which comparative datasets were shaped by paleobiome structure (*w*_B_ > 0) using Bayes factors, provided the strength of the effect was at least moderately strong, even though *w*_B_ is difficult to estimate precisely (Fig. 4). We then fitted our new model to estimate ancestral biomes and regions for *Viburnum*. In discussing our results, we focus on two principal aspects of our study: first, an interpretation of our empirical findings in *Viburnum* and how these may inform other studies seeking to estimate ancestral biomes or regions; and, second, an examination of the model’s assumptions and properties, and how the model’s realism may be improved in future work.

### Biome shifts in Viburnum

*Viburnum* first diversified in the Paleocene and Eocene (66–34Ma), a period when boreotropical forests dominated and connected the northern continents (Wolfe 1985; Graham 2011; Willis and McElwain 2014). Cold temperate forests that experienced long freezing periods were globally rare until after the Oligocene (<34 Ma). Although we inferred an East Asian origin regardless of what biome structure model was assumed, ancestral biome estimates under the three structure models differed in important ways. In the *Paleobiome* analysis, the ancestral biome of the crown node was probably warm temperate (*p* = 0.88) and possibly tropical (*p* = 0.09), and a cold temperate origin could decisively be ruled out (*p* < 0.05; Fig. 5A). When we assumed that biome structure had always resembled today’s structure (*Modern Biome*), the crown node support changed, instead favoring a cold temperate (*p* = 0.67) or possibly a warm temperate (*p* = 0.31) origin (Fig. 5B). The *Null Biome* reconstruction also recovered a warm (*p* = 0.52) or cold (*p* = 0.45) temperate origin, despite the fact that the *Null Biome* inference assumed that all biomes are present in all regions at all times (Fig. 5C). Mismatches between lineage biome affinities and regionally available biomes were highest among pre-Oligocene lineages (>34 Ma). Though cold temperate lineages remained in low proportions (∼5%) until the Oligocene under the *Paleobiome* analysis (Fig. 6A), the *Modern/Null Biome* analyses maintained high proportions of cold temperate lineages in East Asia (> 30%) and North America (7%) in the Eocene (Fig. 6B,C). Over 53% and 47% of pre-Oligocene branches bore mismatched biomes under the *Modern* and *Null Biome* analyses, respectively, but only 6% of those branch lengths were mismatched with biomes under the *Paleobiome* model (Fig. 6D–F). Because of the global rarity of the cold temperate biome during the period of early *Viburnum* evolution, we favored the warm temperate or tropical origin of *Viburnum* under the *Paleobiome* analysis.

Yet, despite stark differences in the *Paleobiome* and *Modern Biome* models, parameter estimates under both conditions found the spatial distribution of biomes to be the primary factor in explaining how viburnums came to live where they do today (*w*_B_ > 0.92, i.e. compatible with the Very Strong condition used in the simulation experiment). Because the ability to estimate ancestral states or to fit evolutionary parameters decays as the evolutionary timescale deepens, we expect that both the *Paleobiome* and *Modern Biome* analyses primarily fit their parameters to phylogenetic patterns of variation pronounced at the shallowest timescales. True to this, model selection by Bayes factors only slightly prefers the *Paleobiome* over the *Modern Biome* structure. All else being equal, however, older *Viburnum* lineages should disperse and biome shift in a manner that is similarly limited by geographical opportunities. The static geographical opportunities assumed under the *Modern Biome* structure induced stationary probabilities that project today’s colder conditions back into the Late Cretaceous, while the dynamic *Paleobiome* structure favored hotter conditions unlike those at present (Figs. 2 and 7). The lesson we take from this is that inferring the fundamental behavior of the process is not always sufficient for estimating ancestral states; inferring if and how that behavior responds to changing historical conditions is also necessary.

We note that an East Asian origin in warm temperate or tropical forests is consistent with several other relevant lines of evidence developed in the study of *Viburnum* evolution, biogeography, and ecology. Previous efforts to reconstruct the ancestral biome of *Viburnum* have weakly favored warm temperate (Spriggs et al. 2015) or cold temperate (Lens et al. 2016) conditions; neither study definitively supported or ruled out a cold temperate origin. Similarly, Edwards et al. (2017) established a relationship between cold temperate conditions and the evolution of deciduousness in *Viburnum*, but could not resolve whether the MRCA was deciduous (cold-adapted) or evergreen (tropical or warm-adapted). Landis et al. (2020) estimated a warm temperate origin of *Viburnum*, with no support for a cold temperate origin, through a combined-evidence tip-dating analysis (Ronquist et al. 2012) that included fossil pollen coded with biome characters to inform the ancestral biome estimates. As a fossil-based estimate, the finding of a non-freezing origin of *Viburnum* cannot be accepted unconditionally; the estimate depends crucially upon the accuracy of biome state assignments to the fossil taxa, and also upon the spatial and temporal biases inherent to fossil deposition and recovery. But, importantly, the fossil-aware biome estimates of Landis et al. (2020) were obtained under the equivalent of our *Null Biome* model, while the fossil-naive estimates in the present study were obtained under the *Paleobiome* model. It is highly satisfying that both studies rule out a cold temperate ancestry for *Viburnum*, and that they do so by leveraging alternative lines of paleobiological evidence: the phylogenetic placement of fossils assigned to particular biomes in one case, and the inferred spatial distribution of biomes through time in the other.

Examining only extant *Viburnum* species, the clade displays considerable variation in both which biomes and which regions lineages occupy. Yet, each region does not contain equal proportions of lineages with affinities to the three biomes. There are several possible causes for this imbalance. In many cases, lineages may simply inhabit regions that lack certain biomes; it is not surprising that there are no extant tropical lineages in North America given that tropical forests have been marginal there since the Oligocene. In other cases, lineages may not have had long enough periods of time for certain biome shifts. For example, all neotropical lineages are adapted to warm temperate (cloud) forests, yet none of them have adapted to the adjacent tropical forest biome. Given the young age of the neotropical radiation, it is possible that there has not been enough time for them to shift into the accessible tropical forests. In this case we can imagine that biological factors (e.g., interactions with other species—competitors, herbivores, etc.—that have long occupied tropical forests) may have played a significant role in limiting this shift (Donoghue and Edwards 2014). In other cases, the imbalance may concern differential rates of speciation or extinction within biomes. For instance, there are relatively few tropical *Viburnum* species given the age and region of origin for the clade and given the age of Asian tropical biomes. If tropical viburnums experienced increased extinction rates (or decreased speciation rates) as they remained in an older biome, that effect would give rise to a pattern of scattered, singular, distantly related, and anciently diverging tropical lineages (depauperons of Donoghue and Sanderson 2015). This is precisely what we see in the case of *V. clemensiae*, *V. amplificatum*, and *V. punctatum* (Spriggs et al. 2015). From analyses under our simple *Paleobiome* model, it appears that temporal, geographical, and ecological influences on rates of character evolution and lineage diversification may all be important factors in explaining why *Viburnum* is distributed as it is across regions and biomes.

Finally, although we question the general validity (often assumed) of stepwise series of events (e.g., trait-first versus climate-first in the evolution of cold tolerance; Edwards et al. 2015), we nevertheless explored how incorporating information on the past distribution of biomes might influence the inference of biome-first versus region-first event series.

Specifically, we asked whether *Viburnum* lineages tended to shift biomes first or disperse to a new region first when radiating through the mesic forests of Eurasia and the New World. Taking the mean proportions across time intervals, we found that when *Viburnum* lineages both disperse into new regions and shift into new biomes, region-first event series (28% of series) are more common than biome-first (19%) series under the *Paleobiome* model.

Alone, this result is difficult to interpret, since the relative number and size of biome and region states will influence what constitutes a biome shift or dispersal event. Using the *Modern* analysis as a point of reference, we find a comparatively neutral relationship, with roughly equal proportions of biome-first (20%) and region-first (21%) series, while under the *Null Biome* analysis the *Paleobiome* relationship is inverted (biome-first, 22%; region-first, 19%). When all regions contain all biomes (*Null Biome*), it makes sense that the ratio of biome-first to region-first series is highest, and that it decreases when the distribution of biomes is not uniform across regions (*Paleobiome* and *Modern Biome*). In the case of *Viburnum*, it appears that several key regional shifts between Eastern Asia and North America occurred a relatively long time ago, when northern latitudes were still primarily covered by warm temperate forests (Fig. 8A). The biome shifts into cold temperate forests occurred later, as cooling climates spread across communities that were already assembled, which is compatible with the ‘lock-step’ hypothesis of Edwards et al. (2017).

Consistent with this scenario, we found that region-first event series do not become the most common series type (over 35%) until the Late Oligocene under the *Paleobiome* model (Fig. 8D). Such region-first event series have also been inferred in several recent analyses, most notably by Gagnon et al. (2019) who found that *Caesalpinia* legumes moved frequently among succulent biomes on different continents, and only later shifted into newly encountered biomes within each continent (Donoghue 2019). From our findings, we suspect that ignoring paleobiome structure may cause the number of region-first transition series to be underestimated. However, it must be borne in our minds that our results may in part reflect the constraint built into our model that simultaneous shifts in biome and region are not allowed (discussed below). In any case, explicitly testing for the effect of paleobiome structure on event order will be important in evaluating patterns of supposed phylogenetic biome conservatism (Crisp et al. 2009).

### Model discussion

Although our model is simple, it is designed with certain statistical features that would allow the model to be applied to diverse datasets beyond *Viburnum*, and to facilitate extensions of the model towards more sophisticated designs. First, we treat many elements in the evolutionary process as model parameters, whose values we estimate from the phylogenetic dataset in question. For example, the *w* parameters control which layers of the paleobiome graphical structure are most relevant to the evolutionary process, and the *y* parameter controls how important weak structural features are when modeling dispersal or biome shift events. Second, the Bayesian modeling framework we chose is ideal for managing complex and parameter-rich hierarchical models (Höhna et al. 2016), allowing for future models to explore the importance of other factors highlighted in the conceptual model of Donoghue and Edwards (2014) — geographical distance (Webb and Ree 2012), region size (Tagliacollo et al. 2015), biome size and shared perimeter (Cardillo et al. 2017), ecological distance (Meseguer et al. 2015), and the effect of biotic interactions on trait and range evolution (Quintero and Landis 2019) — by introducing new parameterizations for computing the time-stratified rate matrices, *Q*(*m*). Our Bayesian framework is also capable of handling sources of uncertainty in the paleobiome graphs, such as uncertainty in the importance of various spatial features or in the age of the appearance of a biome within a region (Landis et al. 2018).

In our application of the model to *Viburnum*, we defined only three biomes and six regions, but the general framework translates to other biogeographical systems with different regions and biomes, provided one can construct an adequate time series of paleobiome graphs. Though our literature-based approach to paleobiome graph construction was somewhat subjective, we found it to be the most integrative way to summarize varied global biome reconstructions, as most individual studies are purely qualitative (Wolfe 1985; Morley 2000; Jetz and Fine 2012; Willis and McElwain 2014; Graham 2011; Graham 2018; but see Kaplan et al. 2003) and based on disparate lines of paleoecological, paleoclimatological, and paleogeological evidence. We believe that our paleobiome graphs for the Northern Hemisphere are sufficiently accurate to show that spatial and temporal variation in the distribution of tropical, warm temperate, and cold temperate forest biomes in space and time can influence how species ranges and biome affinities evolve over time. Nonetheless, future studies should explore more quantitative approaches to defining paleobiome structures for use with the time-stratified regional biome shift model.

Our simple model of regional biome shifts lacks several desired features. Perhaps most importantly, lineages in our model may only occupy a single region and a single biome at a time when, in reality species may occupy multiple biomes and/or regions (e.g., *Liquidambar styraciflua* is a tree that thrives both in temperate deciduous forests of eastern North America and in the cloud forests of Mexico). On paper, it is straightforward to extend the concepts of this model to standard multi-character models, such as the Dispersal-Extinction-Cladogenesis model (Ree et al. 2005; Matzke 2014; Sukumaran et al. 2015). As a DEC model variant, lineages would be capable of gaining affinities with any biomes available within their range. For example, for *M* biomes and *N* regions, there are on the order of 2^ÑÖÜ^ combinations of presences and absences across biomes and regions, and on the order of 2^ÑÜ^ combinations if region-specific biome occupancies are considered.

Computationally, this creates a vast number of viable state combinations, many of which cannot be eliminated from the state space (Webb and Ree 2012). Such a large state space will hinder standard likelihood-based inference procedures for discrete biogeography (Ree and Sanmartín 2009), though recent methodological advances addressing this problem should prove useful (Landis et al. 2013; Quintero and Landis 2019).

Geographical state-dependent diversification (GeoSSE) models may also be interfaced with our model. Incorporating the effect of biome availability on the extinction rate would, at a minimum, be a very important contribution towards explaining patterns of extant diversity. For example, tropical biomes have declined in dominance since the Paleogene, and many ancient *Viburnum* lineages may have since gone extinct in the tropics, perhaps owing to biotic interactions (the dying embers hypothesis of Spriggs et al. 2015). In this sense, we expect that our model will overestimate how long a lineage may persist in a region that lacks the appropriate biome, since our model does not threaten ill-adapted species with higher extinction rates. Efforts to extend GeoSSE models in this manner will face similar, if not more severe, challenges to those encountered in the DEC framework, both in terms of computational limits and numbers of parameters (Beaulieu and O’Meara 2016; Caetano et al. 2018).

If diversification rates vary conditionally on a lineage’s biome-region state, then so should the underlying divergence time estimates. At a minimum, one should jointly estimate divergence times and diversification dynamics to correctly propagate uncertainty in phylogenetic estimates through to ancestral state estimates (Höhna et al. 2019). Beyond that, paleogeographically structured models of biogeography have been shown to be useful for estimating divergence times (Landis 2017; Landis et al. 2018). Paleoecological models, such as our *Paleobiome* model, could be useful in some cases, perhaps for dating clades where some degree of phylogenetic niche conservatism can be safely assumed (Wiens and Donoghue (2004); Crisp et al. (2009); but see Donoghue and Edwards (2014) for potential pitfalls with this approach). For instance, Baldwin and Sanderson (1998) hypothesized that continental tarweeds (*Madiinae*, Asteraceae) radiated within the seasonally dry California Florisitic Province only after Miocene aridification created the province. Baldwin and Sanderson then translated this relationship between biome age and biome affinity to date the maximum crown age of tarweeds, and thus date the maximum crown age of a notable radiation nested within the tarweeds, the Hawaiian silversword alliance. In the future, rather than calibrating the age of tarweeds by asserting a paleoecological hypothesis, it would be possible to use our biome shift model to measure the probability of the dry radiation scenario against competing scenarios, thereby dating the tarweeds (or other clades) based on what ecological opportunities they made use of in different areas and at different times (Baldwin and Sanderson 1998; Landis 2017; Landis et al. 2018).

Finally, although we have compared inferences of event series under several biome structure models, and have argued that paleobiome models can influence such inferences, we caution that event series themselves may not be accurate descriptors of some relevant evolutionary scenarios. Notably, we reiterate that our model does not allow for simultaneous shifts in both biome and biogeographic region. In other words, all biome shifts are restricted to occur within a biogeographic region. This makes several important assumptions about the processes involved in biome shifting. The first is that movements between biomes are non-trivial for organisms, and require a period of time that allows a founding population to adapt to its new environment. If this is true, we see two scenarios that would likely describe the vast majority of biome shifts. First, if two biomes are adjacent to one another, repeated propagule dispersal from the ancestral to the novel biome could allow for gradual adaptation to a new habitat. Second, biome shifts could occur “in-situ”, as one major vegetation type gradually transforms into another, in place, as climate change progresses (“lock-step” model of Edwards et al. 2017). Both of these scenarios would occur *within* biogeographical regions, and we think they probably have accounted for biome shifts in *Viburnum*, as well as many other lineages. However, it is also possible that a shift into a new biome could occur *during* a transition from one region into another. For example, a warm temperate-adapted lineage might adapt to colder forests during range expansion through Beringia, or there might be long-distance dispersal of an organism into newly encountered cold temperate forests to which it was pre-adapted.

Furthermore, it is important to note that our restriction of biome shifts to occur within biogeographical regions does not completely preclude long-distance dispersal biome shifts within regions; nothing in our model ensures that ancestral and novel biomes are adjacent to one another within a given region, especially given that the regions often used are very large and environmentally heterogeneous. Such considerations highlight that the model we have presented here is simplistic in some of its basic assumptions. We view it as a start in the right direction, and look forward to extensions that will allow us to test a variety of more nuanced hypotheses.

## CONCLUSION

The potential for a lineage to adapt to new biomes depends in part on the geographical opportunities that the lineage encountered in time and space. In the case of *Viburnum*, we have shown that differing assumptions about the past distribution of biomes can have a significant impact on ancestral biome estimates. And, when we integrate information about the changing distribution of biomes through time, we favor an origin of *Viburnum* in warm temperate or tropical forests, and confidently rule out an origin in cold temperate forests. The confluence of this line of evidence with our analyses based instead on fossil biome assignments (Landis et al. 2020) provides much greater confidence in a result that orients our entire understanding of the direction of evolution and ecological diversification in this clade.

More generally, we hope that our analyses will motivate biogeographers who wish to estimate ancestral biomes to account for variation in the spatial distribution of biomes through time. We also caution that phylogenetic estimates of ancestral biome affinities derived entirely from extant taxa and biomes may be misleading, particularly for older lineages, even should standard statistical diagnostics indicate that the inference model fits the data relatively well. While we have achieved some conceptual understanding of the interplay between biome shifts in lineages and biome distributions over time, many theoretical and statistical problems must still be solved for us to fully appreciate the significance of changing biome availability in generating Earth’s biodiversity. In presenting our simple model, we hope to provoke further inquiry into how life diversified throughout the biomes of an ever-changing planet.

## FUNDING

This work was supported by the NSF Postdoctoral Fellowship (DBI-1612153) to MJL and a Gaylord Donnelley Environmental Fellowship to MJL through the Yale Institute of Biospheric Studies. Our research on *Viburnum* has been funded through a series of NSF awards to MJD and EJE (most recently, DEB-1557059 and DEB-1753504), and in part through the Division of Botany of the Yale Peabody Museum of Natural History.

## ACKNOWLEDGEMENTS

We thank Bryan Carstens, Mark Holder, and two anonymous reviewers for their thoughtful comments. Ignacio Quintero, Nate Upham, David Polly, and Tracy Heath provided useful feedback that helped us frame the context of the research. We also wish to thank Paul Valdes, who provided us with various BIOME4 reconstructions. Finally, we express gratitude to the Donoghue, Edwards, and Near lab groups for supporting the development of this work since its inception.

## Notes

### Competing Interest Statement

The authors have declared no competing interest.

