## Supplemental Information for "Modeling phylogenetic biome shifts on a planet with a past"

***Supplement 1:** Computing the value of the rate matrix, $Q(m)$*

We provide an example for computing $Q(m)$ for a simple toy system with two biomes (A and B) and two regions (X and Y). Rate matrices for other time intervals are computed similarly for all values of $m$, so we focus only on the first time slice ($m = 1$). In our example, we assume the availability/connectivity of biomes and regions at time $m = 1$ is represented by the adjacency matrix for geography,

$$A_G(1) = \begin{matrix} X \\ Y \end{matrix} \begin{pmatrix} 1 & y \\ y & 1 \end{pmatrix},$$

for biome A,

$$A_A(1) = \begin{matrix} X \\ Y \end{matrix} \begin{pmatrix} 1 & y \\ y & y \end{pmatrix},$$

and for biome B,

$$A_B(1) = \begin{matrix} X \\ Y \end{matrix} \begin{pmatrix} 0 & 0 \\ 0 & 1 \end{pmatrix},$$

where availability (diagonal) or connectivity (off-diagonal) is coded as strong (1), weak ($0 < y < 1$; estimated), or marginal (0). These adjacency matrices are then used to (potentially) rescale the dispersal rates among regions and the biome shift rates among biomes for three rate matrices: for the geographically uninformative rate matrix,

$$Q_U = \begin{matrix} AX \\ AY \\ BX \\ BY \end{matrix} \begin{pmatrix} - & \delta_{XY} & \beta_{AB} & 0 \\ \delta_{YX} & - & 0 & \beta_{AB} \\ \beta_{BA} & 0 & - & \delta_{XY} \\ 0 & \beta_{BA} & \delta_{YX} & - \end{pmatrix},$$

the geography-dependent rate matrix,

$$Q_G(1) = \begin{matrix} AX \\ AY \\ BX \\ BY \end{matrix} \begin{pmatrix} - & [A_G(1)]_{XY}\delta_{XY} & \beta_{AB} & 0 \\ [A_G(1)]_{YX}\delta_{YX} & - & 0 & \beta_{AB} \\ \beta_{BA} & 0 & - & [A_G(1)]_{XY}\delta_{XY} \\ 0 & \beta_{BA} & [A_G(1)]_{YX}\delta_{YX} & - \end{pmatrix}$$

$$= \begin{matrix} AX \\ AY \\ BX \\ BY \end{matrix} \begin{pmatrix} - & y\delta_{XY} & \beta_{AB} & 0 \\ y\delta_{YX} & - & 0 & \beta_{AB} \\ \beta_{BA} & 0 & - & y\delta_{XY} \\ 0 & \beta_{BA} & y\delta_{YX} & - \end{pmatrix},$$

and the biome-dependent rate matrix,

$$Q_B(1) = \begin{matrix} AX \\ AY \\ BX \\ BY \end{matrix} \begin{pmatrix} - & [A_A(1)]_{XY}\delta_{XY} & [A_B(1)]_{XX}\beta_{AB} & 0 \\ [A_A(1)]_{YX}\delta_{YX} & - & 0 & [A_B(1)]_{YY}\beta_{AB} \\ [A_A(1)]_{XX}\beta_{BA} & 0 & - & [A_B(1)]_{YX}\delta_{XY} \\ 0 & [A_A(1)]_{YY}\beta_{BA} & [A_B(1)]_{YX}\delta_{YX} & - \end{pmatrix}$$

$$= \begin{matrix} AX \\ AY \\ BX \\ BY \end{matrix} \begin{pmatrix} - & y\delta_{XY} & 0\beta_{AB} & 0 \\ y\delta_{YX} & - & 0 & 1\beta_{AB} \\ 1\beta_{BA} & 0 & - & 0\delta_{XY} \\ 0 & y\beta_{BA} & 0\delta_{YX} & - \end{pmatrix}.$$

The value of the rate matrix for the regional biome shift process for a given time slice, $Q(m)$, is then determined as by weighted mixture of these rate matrices. If we suppose that $w = (w_U, w_G, w_B) = (0.1, 0.2, 0.7)$, then we have

$$Q(1) = 0.1Q_U \times 0.2Q_G(1) \times 0.7Q_B(1).$$

To assign exact rate values to $Q(1)$, we set $\delta_{XY} = \delta_{YX} = 0.6$, $\beta_{AB} = \beta_{BA} = 0.4$, and $y = 0.3$, producing four dispersal event rate values

$$[Q(1)]_{(AX),(AY)} = 0.6 \times \big((0.1)(1.0) + (0.2)(0.3) + (0.7)(0.3)\big) = 0.222$$

$$[Q(1)]_{(AY),(AX)} = 0.6 \times \big((0.1)(1.0) + (0.2)(0.3) + (0.7)(0.3)\big) = 0.222$$

$$[Q(1)]_{(BX),(BY)} = 0.6 \times \big((0.1)(1.0) + (0.2)(0.3) + (0.7)(0.0)\big) = 0.096$$

$$[Q(1)]_{(BY),(BX)} = 0.6 \times \big((0.1)(1.0) + (0.2)(0.3) + (0.7)(0.0)\big) = 0.096$$

and four biome shift rates values

$$[Q(1)]_{(AX),(BX)} = 0.4 \times \big((0.1)(1.0) + (0.2)(1.0) + (0.7)(0.0)\big) = 0.120$$

$$[Q(1)]_{(BX),(AX)} = 0.4 \times \big((0.1)(1.0) + (0.2)(1.0) + (0.7)(1.0)\big) = 0.400$$

$$[Q(1)]_{(AY),(BY)} = 0.4 \times \big((0.1)(1.0) + (0.2)(1.0) + (0.7)(1.0)\big) = 0.400$$

$$[Q(1)]_{(BY),(AY)} = 0.4 \times \big((0.1)(1.0) + (0.2)(1.0) + (0.7)(0.3)\big) = 0.204$$

with all remaining non-diagonal elements being zero, and each diagonal element equaling the negative sum of the rates departing that row's biome-region state. When fitting the time-stratified regional biome shift model to phylogenetic datasets, we populate the rate matrices for each time interval in a similar manner, $Q = \big(Q(1), Q(2), \ldots, Q(M)\big)$, and further multiply each rate matrix by the global scaling factor, $\mu$, to control the overall rate of the process. Note that dispersal rates between regions X and Y are symmetrical, while shifts between biomes asymmetrical, due to differences in the availability in biomes A and B within regions X and Y.

### ***Supplement 2:*** *Sensitivity analysis of biome structure models*

Biased models may predispose inferences towards certain results, despite the presence of data that would otherwise support alternative results. In our case, it is possible that inferences drawn from our posterior distribution of stochastic mappings (e.g. how 'biome match' proportions differ before and after the Oligocene; Fig. 6) are not driven by information in the data through the likelihood function, but rather that the signal is determined largely by constraints imposed by the *Paleobiome*, *Modern Biome*, and *Null Biome* structure models. We wished to address this concern. To do so, we viewed the biome structure models as empirically structured priors, which allowed us to ask whether those priors have outsized influence on posterior estimates. Put another way, does our choice of empirically structured prior induce biased results?

To assess whether this was an issue, we performed a sensitivity analysis by fitting the *Paleobiome*, *Modern Biome*, and *Null Biome* models to the *Viburnum* dataset in the same manner as described in the main text, with three notable changes. First, we forced the model likelihood function to return the value of 1 for all prior settings. In a Bayesian context, this will cause the posterior distribution to be identical to the prior. Even though the prior is technically data-independent, we nonetheless generated stochastic mappings that conditioned on the biome-region states at the tips of the phylogeny when sampling model parameters from the prior (not the posterior). These stochastic mappings therefore represent a sample of evolutionary histories that are compatible with the comparative dataset under an assumed biome structure model (*Paleobiome*, *Modern Biome*, and *Null Biome*) for parameter values as supported by our prior. Standard stochastic mapping algorithms are typically applied using model parameters that explain the data well (e.g. using maximum likelihood estimators or posterior samples for model parameters). Much of prior parameter space, however, explains a particular dataset extremely poorly. We found that our stochastic mapping algorithm was numerically unstable for some of our most extreme rates (e.g. when $\mu$ equaled $10^{-4}$ or $10^{1}$ events/Myr). To improve algorithmic stability, we made our second modification to the original analysis settings, by using a more conservative prior on the global event rate for biome-region state transitions, substituting the original prior of $\mu \sim$ Loguniform($10^{-4}$, $10^{1}$) with $\mu \sim$ Loguniform($10^{-3}$, $10^{0}$). Posterior estimates under the original prior were generally in the range of $10^{-2}$ to $10^{-1}$ and exclude values as extreme as $10^{-4}$ or $10^{1}$, and both priors share an expectation of $10^{-1}$, so we believe this substitution is wholly appropriate given the unusual nature of the problem we are attempting to address. Finally, our sensitivity analyses differed in that we ran MCMC for fewer iterations to collect the same number of samples, since it is generally easier to sample from the prior than it is to do so from the posterior. In addition to generating the three induced prior distributions of stochastic mappings, we also generated the induced prior distribution of root state frequencies.

Our sensitivity analyses show that regardless of which biome structure model we assumed, all reconstructions produced nearly identical lineage state proportions through time (Fig S1A−C) and nearly identical lineage-biome match proportions through time (Fig S1D−F). In general, posterior estimates for lineage-state proportions supported only an East Asian origin of *Viburnum* (Fig. 6A−C), whereas our prior-based estimates awarded diffuse support to a broad range of alternative regions of origin (Fig. 6D−F). Conflicting posterior estimates of a tropical, warm temperate, or cold temperate origin of *Viburnum* that were contingent on which biome structure model was analyzed (Fig. 6A−C) is completely erased under the prior-based inference (Fig. S1A−C). Lineage-biome match proportions under the prior lingered around 65% until the end of the Oligocene, after which they rose to ~85%, which are most similar to the posterior estimates under the *Modern* and *Null* biomes (Fig. 6E,F). By contrast, posterior estimates under the *Paleobiome* model (Fig. 6D) inferred high proportions of lineage-biome matches (>95%) over all time intervals.

Prior distributions of root state frequencies were generally insensitive to which biome structure model was assumed (Fig. *S*2). The *Paleobiome, Modern Biome*, and *Null Biome* structures all have median state frequencies that are close $1/18 \approx 0.056$, i.e. the value one expects if all 18 states had equal prior probability. Highest posterior densities are also similar across biome structure models and biome-region states. In contrast, posterior root state frequency estimates (Fig. 7) departed significantly from the value 0.056, and in ways that reflect the given biome structure model. For example, the small posterior probabilities for the Cold+SEAs and Cold+SAm root states do not include the value 0.056 in their 95% highest posterior densities (HPDs). In contrast, the *Null Biome* prior and posterior density have fairly similar medians, although their HPDs differ.

We did not find compelling support that our posterior estimates for *Viburnum*, especially for those results under the *Paleobiome* model, are due to inherent and overwhelming biases in the underlying biome structure models. Rather, posterior inferences under alternative biome structure models differed in large part because of how each model fit its parameters to the datasets through the model (i.e. through the likelihood function). Because these results will not hold for any comparative dataset or for any biome structure model imaginable, future researchers wishing to use our biome shift model to test hypotheses in other empirical systems are advised to perform their own sensitivity analyses.

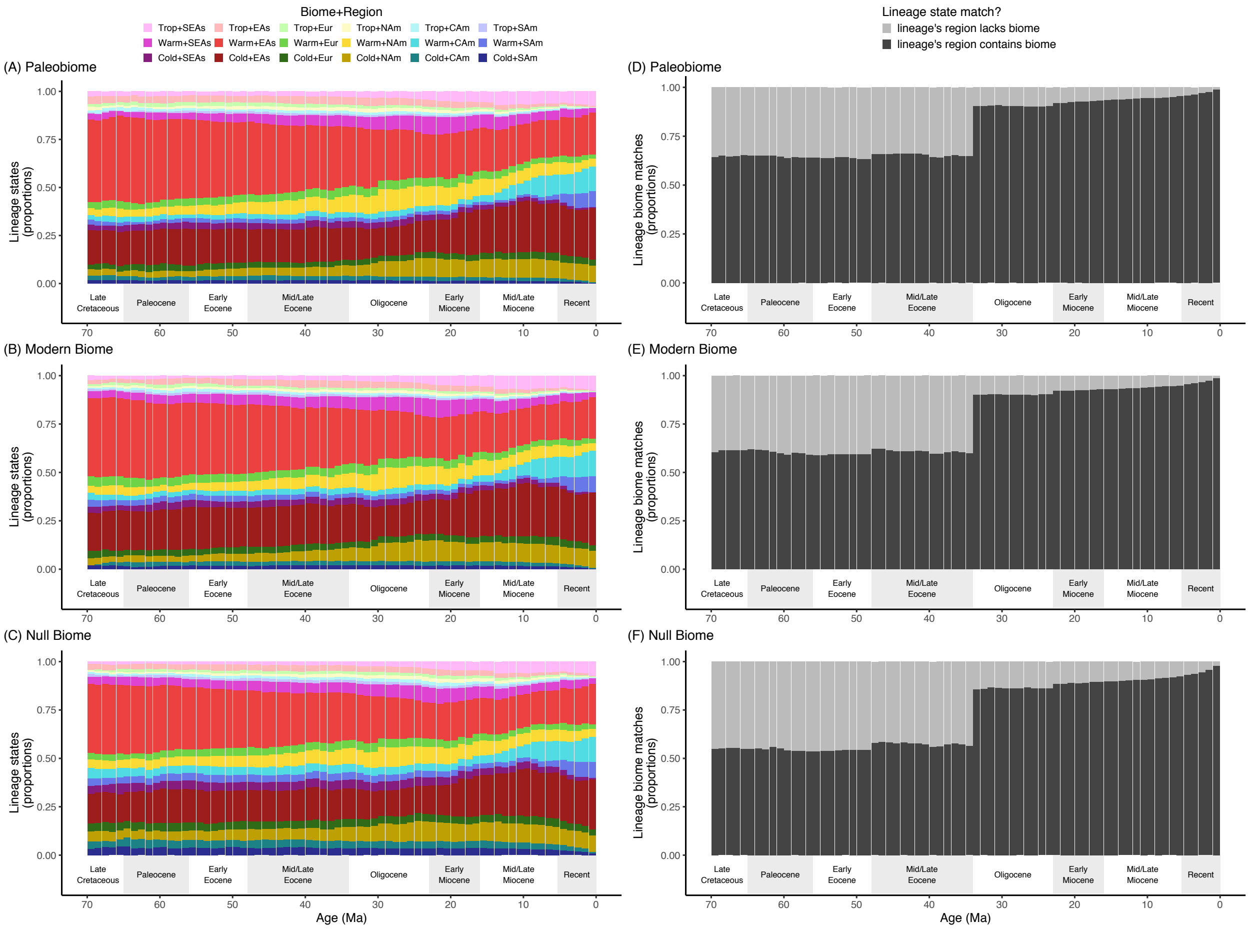


**Figure S1.** Ancestral proportions of lineage state frequencies through time for *Viburnum* as estimated under the prior (Supplement 2). The left column (A–C) shows the lineages biome-region states, where regions differ by color and biomes differ by shading (see legend). Proportions of reconstructed lineages in each biome-region state are shown for estimates under the *Paleobiome* (A), *Modern Biome* (B), and *Null Biome* (C) settings. The right column (D–F) shows the proportion of lineages with biome states that match (dark) or mismatch (light) the non-marginal biomes that are locally accessible given any lineage's location, as defined under the Paleobiome structure (see main text for details). Proportions of reconstructed lineages with biome match and mismatch scores are shown for estimates under the *Paleobiome* (D), *Modern Biome* (E), and *Null Biome* (F) settings.

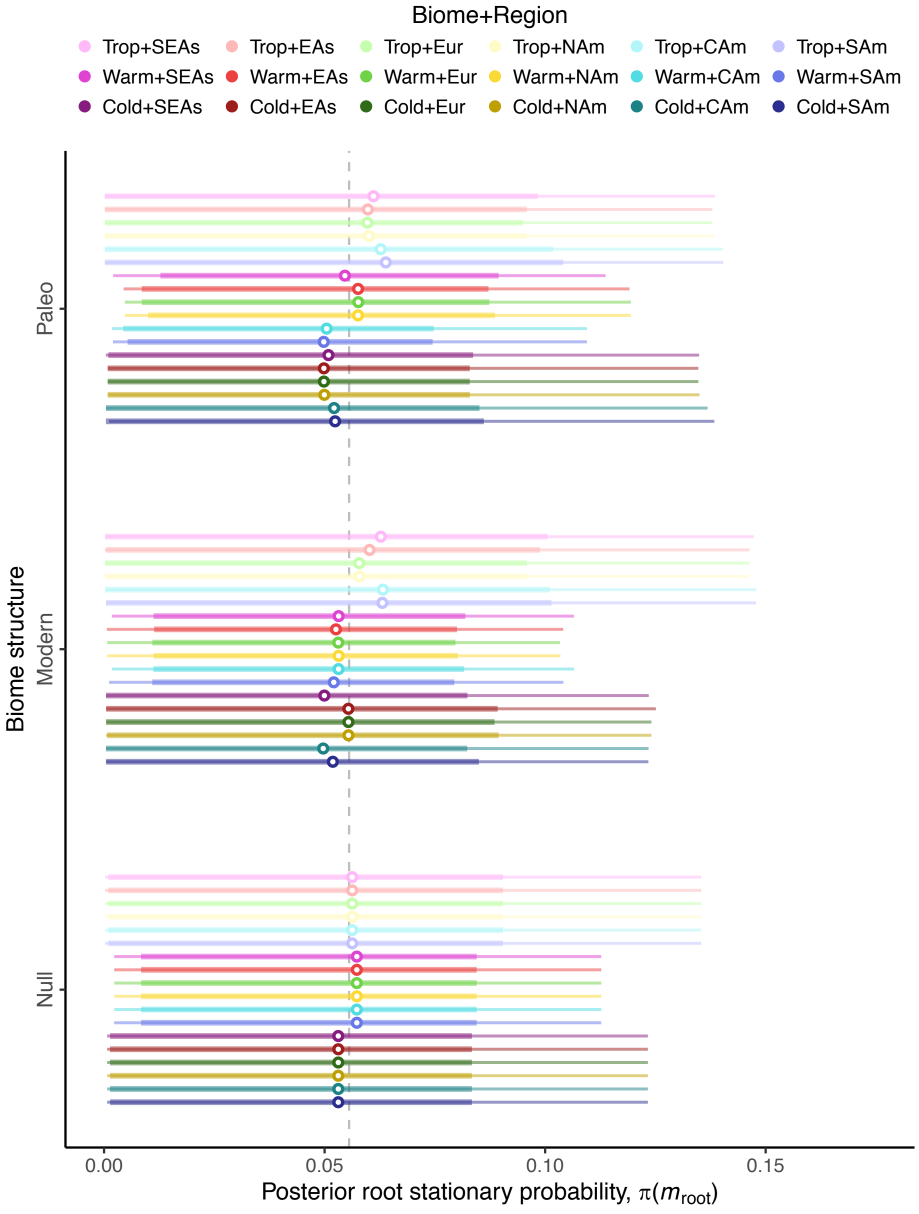


**Figure S2.** Stationary probabilities for the *Viburnum* root state during the Late Cretaceous under the prior. Prior stationary probabilities for $\pi(m_{root})$ are given for each biome structure model (grouped rows) and for each biome-region state (colors) as posterior means (points) and credible intervals (HPD80, thick lines; HPD95, thin lines).
